# An uncommon phosphorylation mode regulates the activity and protein-interactions of N-acetylglucosamine kinase

**DOI:** 10.1101/2024.02.23.581760

**Authors:** Arif Celik, Ida Beyer, Dorothea Fiedler

## Abstract

While the function of protein phosphorylation in eukaryotic cell signaling is well established, the role of a closely related modification, protein pyrophosphorylation, is just starting to surface. A recent study has identified several targets of endogenous protein pyrophosphorylation in mammalian cell lines, including N-acetylglucosamine kinase (NAGK). Here, a detailed functional analysis of NAGK phosphorylation and pyrophosphorylation on serine 76 (S76) has been conducted. This analysis was enabled by using amber codon suppression to obtain phosphorylated pS76-NAGK, which was subsequently converted to site-specifically pyrophosphorylated NAGK (ppS76-NAGK) with a phosphorimidazolide regent. A significant reduction in GlcNAc kinase activity was observed upon phosphorylation, and near-complete inactivation upon pyrophosphorylation. The formation of ppS76-NAGK proceeded *via* an ATP-dependent autocatalytic process, and once formed, ppS76-NAGK displayed notable stability towards dephosphorylation in mammalian cell lysates. Proteomic examination unveiled a distinct set of protein-protein interactions for ppS76-NAGK, suggesting an alternative function, independent of its kinase activity. Overall, a significant regulatory role of pyrophosphorylation on NAGK activity was uncovered, providing a strong incentive to investigate the influence of this unusual phosphorylation mode on other kinases.

## Introduction

The covalent post-translational modification (PTM) of proteins is a widely used mechanism to regulate protein structure and function within cells, and an impressive array of PTMs has been documented to date.^2^ Protein phosphorylation is one of the most prevalent modifications and plays a central role in cellular signal transduction processes. It has been estimated that 30-65% of the human proteome are phosphorylated at some point in time, however, the functional relevance for the majority of phosphorylation sites remains unclear.^3-4^

To interrogate the impact of phosphorylation on protein structure and activity, access to stoichiometrically phosphorylated proteins is of great benefit. Chemical tools offer a precise means to synthesize stoichiometrically phosphorylated proteins. Protein semi-synthesis, a combination of solid-phase peptide synthesis (SPPS) to include the desired phosphorylation site, and protein expression of protein fragments, is a powerful technique to obtain stoichiometrically modified proteins and has been widely applied.^8—8^ In addition, the stoichiometric and site-specific incorporation of unnatural amino acids *via* genetic code expansions has recently been advanced to include the incorporation of phosphoserine, phosphothreonine, and phosphotyrosine.^9—12^

Adding to the complexity of phosphorylation-based signaling, a novel, non-enzymatic modification, termed protein pyrophosphorylation, was reported in 2007.ref This modification is thought to be mediated by inositol pyrophosphate messenger (PP-InsPs),which putatively transfer their β-phosphoryl group to pre-phosphorylated protein substrates, resulting in the formation of a diphosphate (pyrophosphate) moiety.^13—14^ Our group recently developed a mass spectrometry approach to identify many endogenous mammalian pyrophosphorylation sites within complex samples.^15^ The pyrophosphorylation sites were typically detected on nuclear and nucleolar proteins, and the sites localized to acidic polyserine stretches with a high degree of disorder.

However, a few pyrophosphoproteins did not fit that picture, among them N-acetylglucosamine kinase (NAGK). In NAGK, the pyrophosphorylation site is in a structurally resolved area of the protein, immediately adjacent to the substrate binding site (Figure 1). NAGK is best known for its participation in the N-acetylglucosamine (GlcNAc) salvage pathway, where it catalyzes the phosphorylation of GlcNAc to produce GlcNAc-6-phosphate.^16—18^ While GlcNAc-6-phosphate is typically synthesized de novo *via* the hexosamine biosynthesis pathway, in nutrient-deprived microenvironments, cellular GlcNAc-6-phosphate concentrations can be maintained at a high level by an increased expression of NAGK and heightened GlcNAc salvage metabolism.^19^

**Figure 1.**
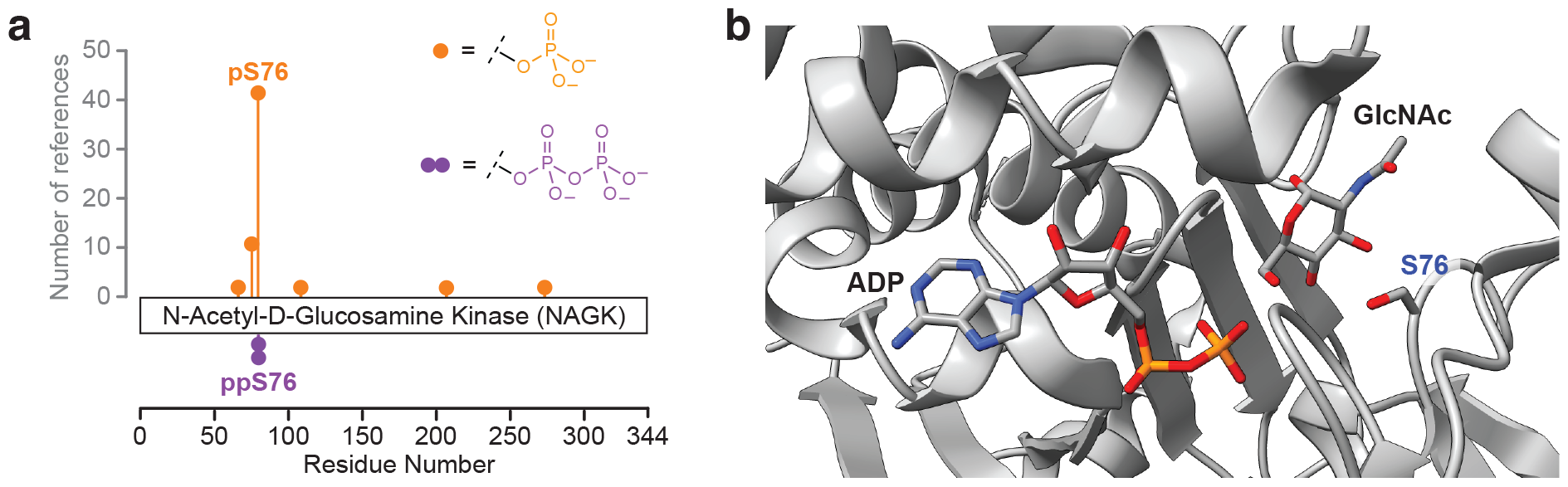
a) Reported phosphorylation sites on NAGK in PhosphoSitePlus by high-throughput MS analysis and previously discovered pyrophosphorylation site S76 reported by Morgan et al. b) Crystal structure of NAGK co-crystalized with ADP and GlcNAc highlighting the location of the ppS76 site. Carbon atom is indicated in grey, Oxygen atom is indicated in red, and phosphorous atom is indicated in orange. PDB: 2CH5 and 2CH6 were merged together.

In addition to its capacity to phosphorylate GlcNAc, NAGK can phosphorylate peptidoglycans, specifically muramyl dipeptide (MDP), to form 6-O-phospho-MDP in a process that triggers pro-inflammatory gene expression.^20—21^ Finally, a few reports have recently ascribed a scaffolding function to NAGK – independent from its catalytic activity.^22—25^

Considering the central role of NAGK in the GlcNAc salvage pathway, and its additional functions, it is important to understand how NAGK activity and function are regulated. While several sites of modification - predominantly phosphorylation - of NAGK have been identified in high-throughput studies, the functional relevance of these modification sites has not been addressed (Figure 1). We were particularly interested in the phosphorylation site on serine 76 (S76), because it is the most commonly detected phosphorylation site on NAGK.^26^ In addition, S76 was recently found to be pyrophosphorylated, and is very close to the substrate binding pocket.^15—16^

To elucidate the influence of phosphorylation and pyrophosphorylation of NAGK at S76, we now report the synthesis of site-specifically phosphorylated and pyrophosphorylated NAGK. Access to these stoichiometrically modified protein samples enabled the detailed characterization of the different phosphorylation modi. Compared to the unmodified protein, the GlcNAc kinase activity of the phosphoprotein was significantly reduced. The pyrophosphoprotein was almost completely inactive. This inactivation appears to be irreversible, as the pyrophosphoryl group on NAGK was resistant to hydrolysis in active cell lysates. Instead, pyrophosphorylation bestows NAGK with the ability to physically interact with a distinct set of proteins, compared to the unmodified protein. Overall, our findings highlight the intricacies of protein regulation, where a subtle change of protein modification can bring about large alterations in behavior, and will motivate us and others to further explore the biochemical and biophysical properties of different pyrophosphorylated kinases.

## Results

### Synthesis of pyrophosphorylated NAGK

To investigate how pyrophosphorylation on S76 may affect the properties and function of NAGK, site-specifically and stoichiometrically modified protein is required. Based on previous work, we sought to apply a combination of amber codon suppression to incorporate phosphoserine, followed by a chemoselective reaction of the phosphoprotein with a photolabile phosphorimidazolide reagent, to obtain the corresponding pyrophosphoprotein.^27—28^ Expression of the phosphoprotein in Escherichia coli proceeded smoothly and yielded good quantities of pS76-NAGK (7.5 mg/L) in high purity (Figure 2, Figures S1 and S2). pS76-NAGK was subsequently treated with biotin-polyethylenglycol-6-triazole-nitrophenylethyl-phosphorimidazolide (biotin-PEG_6_-Tz-NPE-phosphorimidazolide) for 18 h at 45 °C in a solvent mixture of DMA:H_2_O (9:1). After refolding, the formation of the derivatized protein, R-ppS76-NAGK, was clearly observed by quadrupole time-of-flight mass spectrometry (Q-TOF-MS). To release the pyrophosphoprotein, the sample was exposed to 365 nm light, and ppS76-NAGK could be isolated without notable formation of any side products (Figure 2). To confirm that the refolding step was suitable to obtain properly folded protein, we subjected wildtype-NAGK (wt-NAGK) to the derivatization/refolding conditions. Circular dichroism (CD) measurements corroborated the successful restoration of protein structure (Figure S3), and kinase activity was unaltered. In sum, the site-specifically modified phosphoprotein pS76-NAGK, and pyrophosphoprotein ppS76-NAGK, could be accessed readily, alongside the unmodified protein, wt-NAGK.

**Figure 2.**
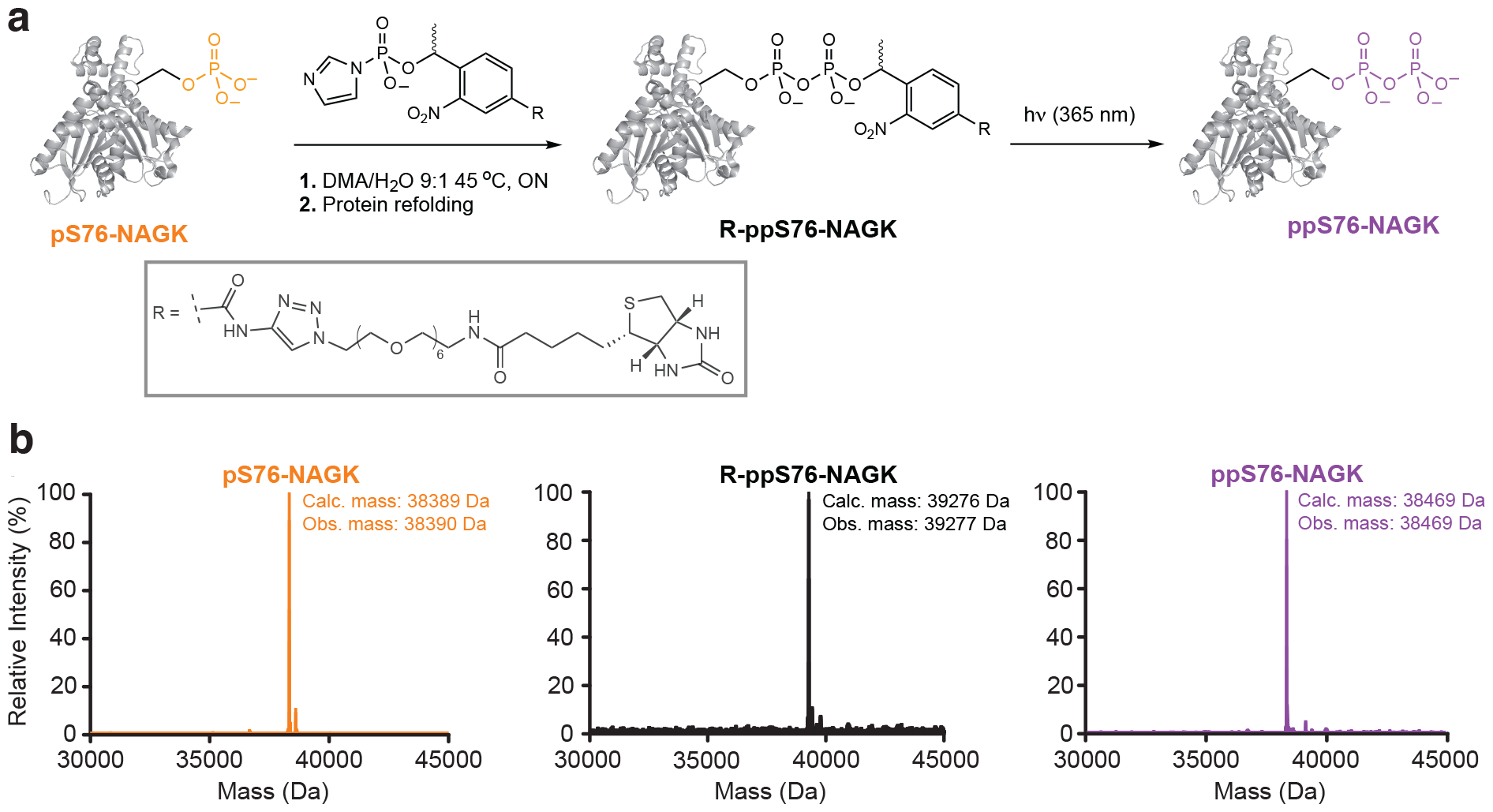
Chemical phosphorylation of pS76-NAGK to provide pyrophosphoprotein ppS76-NAGK. pS76-NAGK is derivatized by using biotin-PEG_6_-Tz-NPE-phosphorimidazolide for the selective modification of the phosphoserine moiety, to yield R-ppS76-NAGK. Subsequent irradiation releases the pyrophosphoprotein ppS76-NAGK. Q-TOF-MS analysis of the intermediates and products is shown below.

### Pyrophosphorylation severely reduces GlcNAc kinase activity

With wt-NAGK, pS76-NAGK, and ppS76-NAGK in hand, we next wanted to evaluate how the different phosphorylation modes influenced the enzymatic activity of NAGK. To do so, a standard kinase assay was set up, using GlcNAc as a substrate and monitoring adenosine-5’-triphosphate (ATP) consumption (Figure 3a).^29^ At low enzyme concentration (1 nM), unphosphorylated wt-NAGK displayed a robust activity of 260 nmol/min/ng. By contrast, pSer76-NAGK showed no activity at 1 nM enzyme concentration. Even with increased enzyme concentration, its overall activity decreased by approximately 200-fold to 1.4 nmol/min/ng (Figure 3a, Figure S4). Pyrophosphorylation on S76 decreased the kinase activity even more, and low conversion only became detectable at an enzyme concentration of 1 µM (Figure 3a). In addition to its GlcNAc-kinase activity, it was recently shown that NAGK can phosphorylate MDP to generate MDP-6-phosphate in response to bacterial infection.^20^ We therefore tested the ability of wt-NAGK, pS76-NAGK, and ppS76-NAGK to utilize MDP as a substrate. Again, wt-NAGK exhibited high activity towards MDP, while the activity of pS76-NAGK was significantly reduced, and ppS76-NAGK showed even lower conversion (Figure 3b). Overall, the kinase activity of NAGK towards GlcNAc and MDP was strongly reduced by the introduction of a single phosphoryl group on S76. Pyrophosphorylation on this side chain caused an even more pronounced effect, making NAGK virtually inactive.

**Figure 3.**
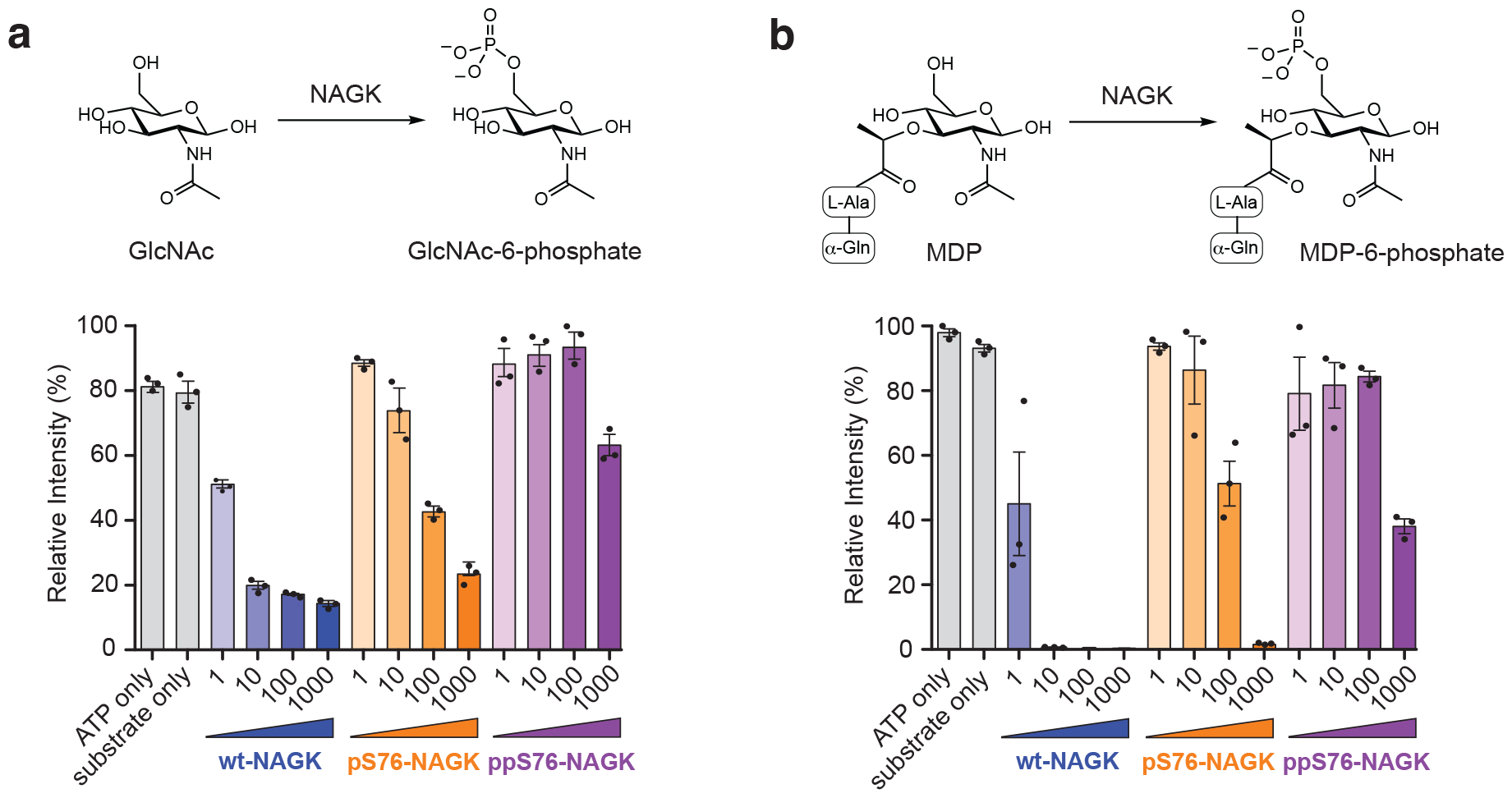
Phosphorylation and pyrophosphorylation of NAGK reduce the enzymatic activity. a) NAGK kinase activity towards GlcNAc as a substrate was performed at 37 °C for 1 h in a) 50 mM HEPES (pH 7.5), 50 mM NaCl, 10 mM MgCl_2_, 70 µM GlcNAc, 100 µM ATP. Concentrations of NAGK are in nM. b) NAGK kinase activity towards MDP as a substrate was performed at 37 °C for 1 h in 50 mM HEPES (pH 7.5), 50 mM NaCl, 10 mM MgCl_2_, 190 µM MDP, 100 µM ATP. Concentrations of NAGK are in nM. wt-NAGK is indicated in blue, pS76-NAGK is indicated in orange, and ppS76-NAGK is indicated in purple. Data presented as mean ± standard error (SE) of three technical replicates.

### Phosphorylation by aurora kinase B is followed by autopyrophosphorylation

Given the pronounced effect of a single phosphorylation/pyrophosphorylation event on NAGK activity, we wondered how these modifications were installed. To date, no biochemical data on protein kinases targeting S76 on NAGK have been reported. Nonetheless, a prior high-throughput proteomic investigation identified this residue as a candidate site for phosphorylation by aurora kinase B (AurB) and cyclin-dependent kinase 1 (CDK1).^30—31^ To validate this potential connection, AurB was incubated with wt-NAGK in the presence of ATP and MgCl_2_. Q-TOF-MS corroborated the addition of the phoshoryl group to NAGK (Figure 4a), and MS/MS analysis confirmed the addition to serine 76 (Figure S4).^32^ Phosphorylation by AurB also notably reduced GlcNAc kinase activity over time (Figure 4b), consistent with our previous observation using recombinantly expressed pS76-NAGK.

**Figure 4.**
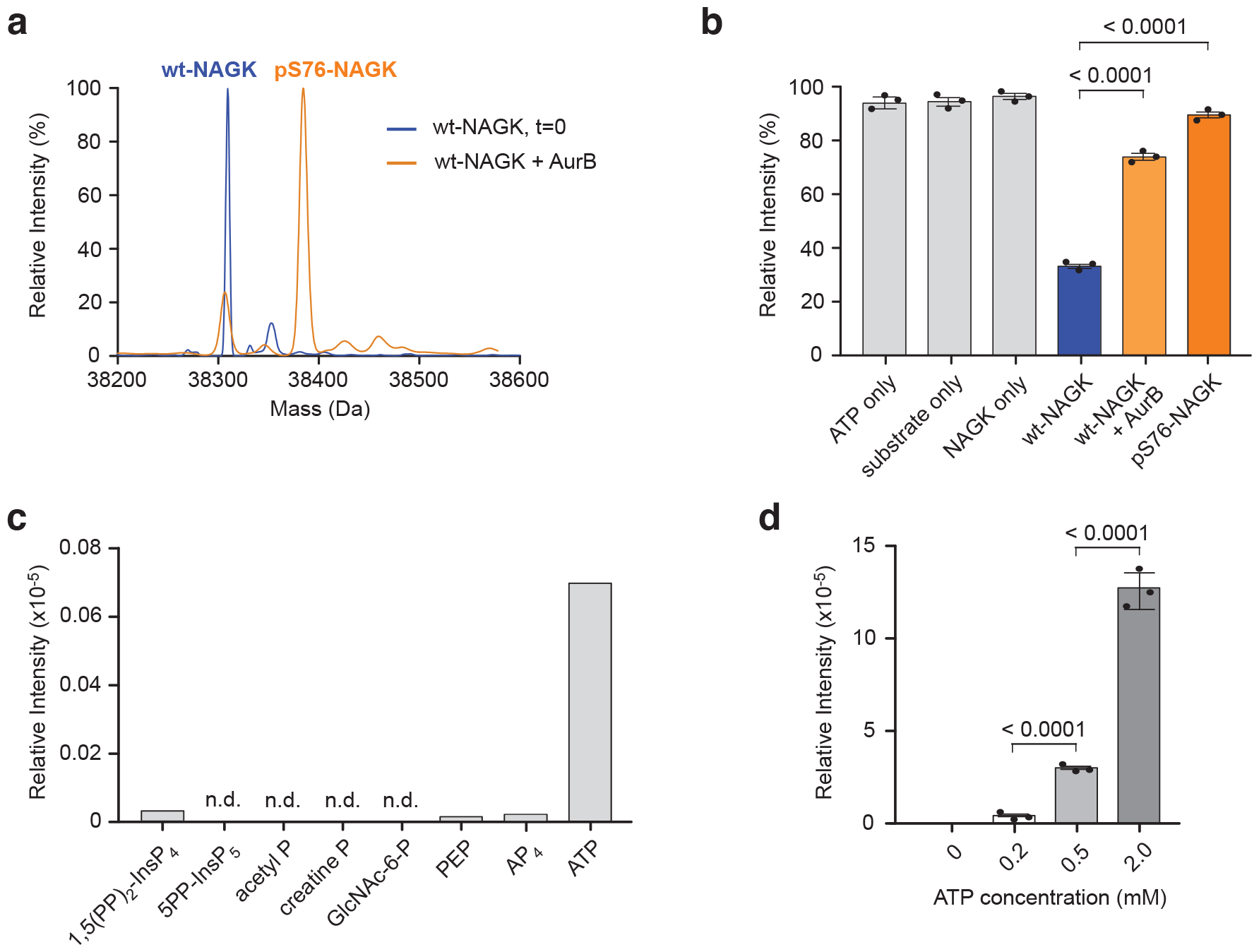
Phosphorylation and pyrophosphorylation of NAGK. a) AurB phosphorylates NAGK. Q-TOF-MS spectra displaying NAGK before (blue) and after (orange) the overnight incubation with AurB. b) Measurement of GlcNAc kinase activity, before and after treatment with AurB. GlcNAc kinase activity is decreased following an overnight incubation with AurB in Tris-Buffer. The activity of pS76-NAGK is shown for comparison. c) Autopyrophosphorylation with different metabolites harboring potential pyrophosphorylation properties analyzed by MS/MS. Assays were performed by incubating at 37 °C overnight in 50 mM HEPES (pH 7.5), 150 mM NaCl, 0.8 mM MgCl_2_, 1 mM DTT, 10 µM pS76-NAGK, and 0.2 mM Phosphoryl-Donor. d) Graph showing the relative quantification of pyrophosphoryation upon increased ATP concentration. Assays were performed by incubating at 37 °C overnight in 50 mM HEPES (pH 7.5), 150 mM NaCl, 0—2 mM MgCl_2_, 1 mM DTT, 10 µM pS76-NAGK, and 0—2 mM ATP. Data presented as mean ± SE of three technical replicates. P-values were determined by unpaired t-test analysis.

Protein pyrophosphorylation is a recently emerging modification and is thought to be mediated non-enzymatically by inositol pyrophosphate metabolites (PP-InsPs).^13—14,33—37^ Therefore, we tested whether ppS76-NAGK would form upon incubation of pS76-NAGK with PP-InsPs, specifically 5-diphospho-inositolpentakisphosphate (5PP-InsP_5_) and 1,5-bisdiphospho-insitoltetrakisphosphate (1,5(PP)_2_-InsP_4_) in the presence of Mg^2+^ ions. Interestingly, neither 5PP-InsP_5_, nor 1,5(PP)_2_-InsP_4_ were capable of generating the pyrophosphoprotein, even after extended reaction times (Figure 4c). While the product of the 1,5-PP-InsP_5_ reaction indicated signs of the mass corresponding to pyrophosphorylation at the MS1 level, MS/MS analysis did not confirm product formation. Because mammalian cells contain various high-energy metabolites that could serve as phosphoryl donors, we subsequently investigated the ability of ATP^38^, adenosine-5’-tetraphosphate (AP_4_)^39^, phosphoenolpyruvate (PEP)^40^, GlcNAc-6-phosphate, creatine phosphate^41^ and acetyl phosphate^42^ to participate in phosphoryl-transfer chemistry. For acetyl phosphate, creatine phosphate, and AP_4_, again, formation of ppS76-NAGK was not detected (Figure 4c). However, upon treatment of pS76-NAGK with 200 μM ATP, a robust signal for the corresponding pyrophosphopeptide became apparent in the MS1 spectrum, and the pyrophosphoryl group could by localized by MS/MS analysis (Figure S6).^30^ Given that no additional enzymes were present in these biochemical reactions, the generation of the pyrophosphoryl moiety appears to be an autocatalytic event. The degree of autopyrophosphorylation should therefore correlate with the ATP concentration. Indeed, the signal for the pyrophosphopeptide increased when ATP concentrations were elevated (Figure 4d). Overall, formation of the pyrophosphate group on ppS76-NAGK is not mediated by PP-InsPs, but is generated in an ATP-dependent, autocatalytic fashion. Considering the close proximity between the ATP binding site and pS76, an intramolecular phosphoryl transfer reaction appears feasible (Figure 1b).^16^ Given the unusual reaction sequence that leads to pyrophosphorylation, the question arises if, and how, this modification can be removed.

### ppS76-NAGK is resistant to dephosphorylation in mammalian cell lysates

To maintain dynamic control over protein function, phosphorylation of proteins is typically reversible, and dephosphorylation is catalyzed by protein phosphatases.^44—45^ Phosphatases that would remove one or two phosphoryl groups from pyrophosphoproteins have not been described to date. To investigate possible dephosphorylation reactions, we prepared cell lysates from HEK293T cells, and first monitored the dephosphorylation of pS76-NAGK by Q-TOF-MS (Figure 5a). High conversion of pS76-NAGK to the dephosphorylated product was observed, validating the activity of the lysate. In contrast, ppS76-NAGK displayed no hydrolysis at all, even after extended incubation times, and using different lewis acidic metals cations in the buffer, demonstrating a high degree of biochemical stability of the pyrophosphoserine group (Figure 5b). The inertness of ppS76-NAGK towards hydrolysis was also maintained in lysates from human colon colorectal carcinoma cells (HCT116) and human pancreas-1 (PANC-1) cells (Figure S8).

**Figure 5:**
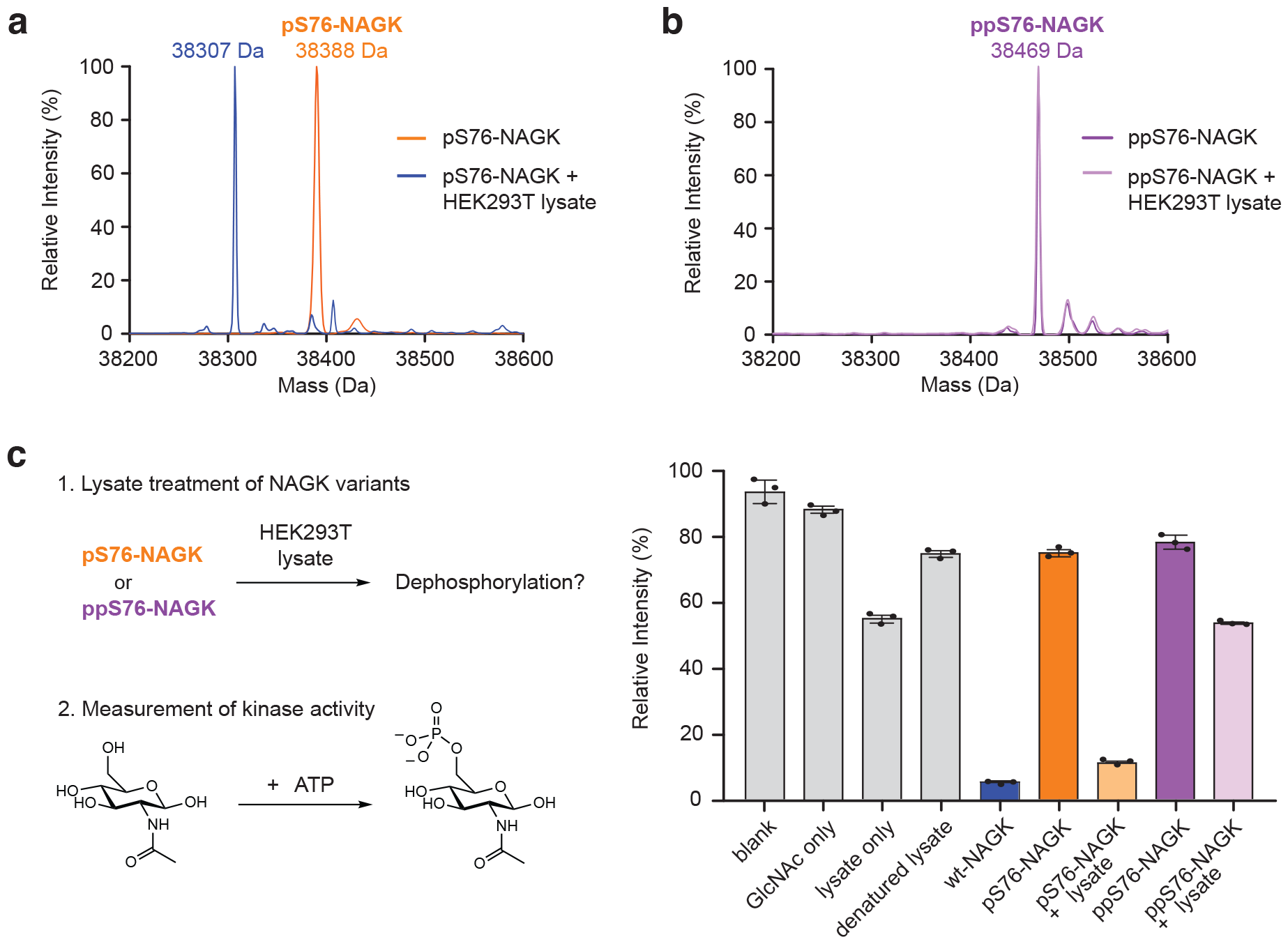
Dephosphorylation of pS76-NAGK and ppS76-NAGK. a) Q-TOF-MS based analysis of HEK293T treated pS76-NAGK clearly shows the dephosphorylation and formation of wt-NAGK. b) Q-TOF-MS based analysis of HEK293T treated ppS76-NAGK was not affected by phosphatases in lysate. c) The presence of potential pyro- and phosphatases in mammalian cells. Biochemical validation of NAGK (wt, pS76, ppS76) after treatment with HEK293T treatment and subsequent GlcNAc kinase activity assay. Subsequent experiments with HCT116 and PANC-1 cell lysate also showed regained activity for pS76-NAGK but not ppS76-NAGK (Figure S7) Data presented as mean ± SE of three technical replicates. The bar graph including t-test is shown in Figure S7.

Since dephosphorylation of NAGK should restore its GlcNAc kinase activity, we next probed this activity upon incubation of pS76-NAGK and ppS76-NAGK with cell lysate. As expected, pS76-NAGK regained kinase activity upon exposure to HEK293T lysates, whereas ppS76-NAGK did not (Figure 5b). The residual kinase activity observed in the ppS76-NAGK sample can be attributed to the presence of ATPases in the lysate, leading to ATP consumption (Figure 5b). These results are in line with previous work using radiolabeled pyrophosphoproteins, where a resistance towards dephosphorylation by common protein phosphatases was reported.^14^ The increased stability of the pyrophosphorylation mark (compared to monophosphorylation) seems to equip the proteins with a more permanent modification, and may potentially play a role in mediating protein-protein interactions.^34,46^

### Pyrophosphorylation influences the protein-protein interactions of NAGK

A few recent examples have shown how protein pyrophosphorylation can either enhance or decrease the affinity of specific protein-protein interactions.^33—38,46^ To investigate how the protein interactome of ppS76-NAGK compared to wt-NAGK, lysates were prepared from HEK293T cells and were incubated with His6-tagged wt-NAGK or ppS76-NAGK. Following immobilization on nickel beads, the supernatants were removed, the protein-complexes were washed and subsequently eluted with imidazole.^47^ Analysis of the eluate by high-resolution mass spectrometry identified many known interactors of NAGK (Figure S8).^48—51^

Next, a volcano plot was generated comparing LFQ values for wt-NAGK versus ppS76-NAGK, to examining how protein-protein interactions were altered by pyrophosphorylation (Figure 6a). Interestingly, several known NAGK interactors (AKAP8, HNRNPH1, HNRNPH2, SCAF8, RBM6, USP15, LNX2, WIPI2, PPIL2, PATL1, DACH1, and YLPM1) no longer associated with pyrophosphorylated NAGK, suggesting that pyrophosphorylation destabilized these interactions (Figure 6a). On the contrary, not a single known NAGK interactor preferentially bound to ppS76-NAGK. Instead, pyrophosphorylation appears to promote a distinct set of protein-protein interactions: 78 proteins were identified that preferentially interacted with ppS76-NAGK, either directly or indirectly (Figure 6a, Table S1).

**Figure 6:**
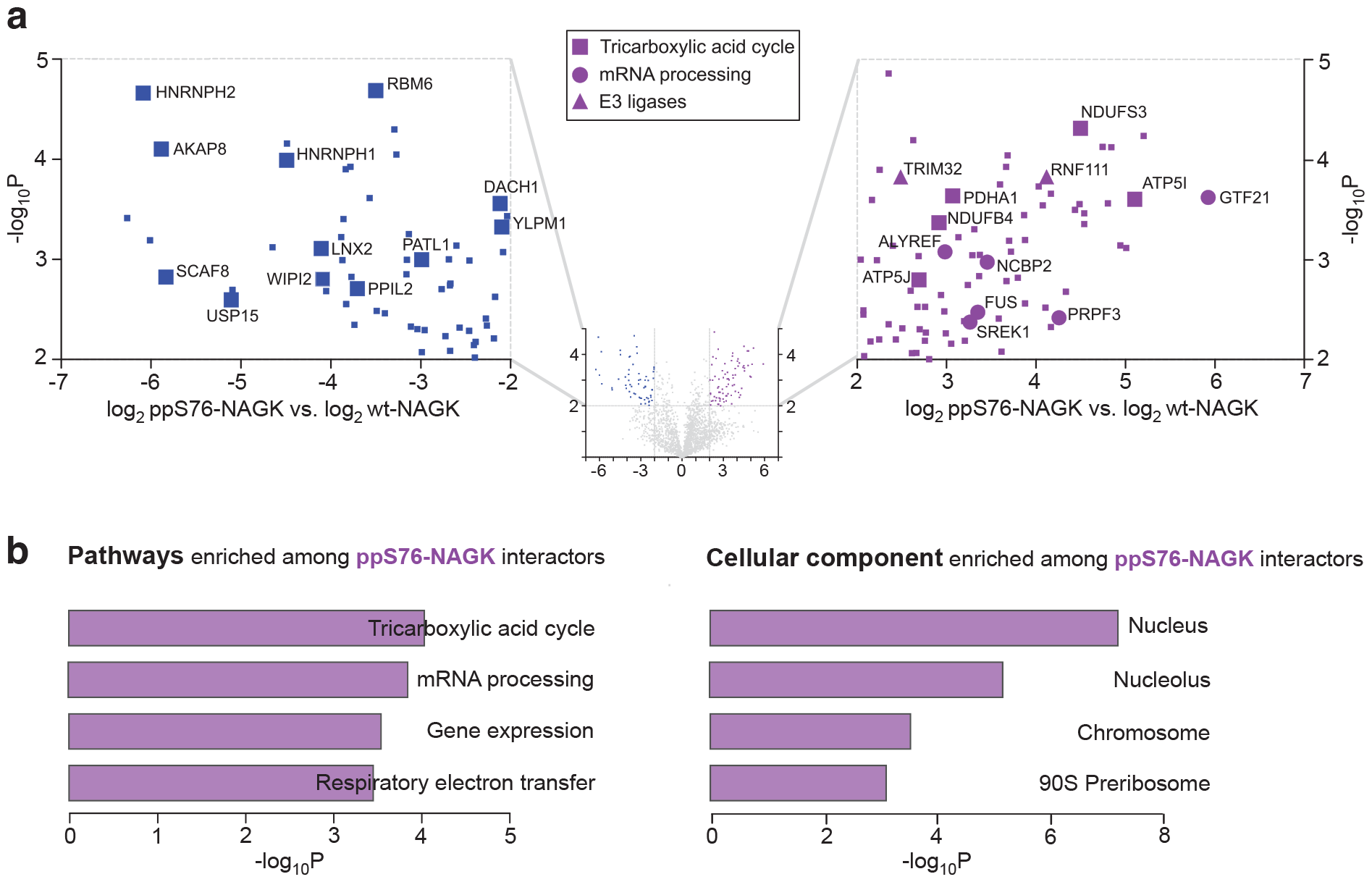
Interactome analysis of pp76-NAGK versus wt-NAGK. a) Volcano plot depicting LFQ values of ppS76-NAGK versus wt-NAGK after a t-test. The x-axis displays the difference of LFQ values on a log2 scale and the y axis shows the −log10p value. Left side illustrates the preferential enrichment with wt-NAGK. Right side illustrates preferential enrichment with ppS76-NAGK. b) Gene ontology analysis of pp76-NAGK interacting proteins.

To associate this list of 78 proteins with functional terms, we used Enrichr for gene ontology analysis (Table S2).^52—53^ This analysis suggests a role of ppS76-NAGK in the tricarboxylic acid (TCA) cycle and respiratory electron transport, as proteins from these pathways (ATP5I, ATP5J, PDHA1, NDUFB4, and NDUFS3) are overrepresented among the ppS76-NAGK interactors (Figure 6b). Additionally, ppS76-NAGK binds to several proteins involved in mRNA processing (GTF2F1, FUS, SREK1, NCBP2, and PRPF3). Consistent with the association with mRNA processing, the nucleus is the cellular component most strongly enriched among ppS76-NAGK interactors (Figure 6b). Finally, we identified two E3 ligases, TRIM32 and RNF111, that preferentially interacted with ppS76-NAGK. Given the biochemical stability of pyrophosphorylation, it is plausible that - instead of dephosphorylation - ubiquitin-mediated degradation may be involved in the turnover of pyrophosphorylated NAGK.^46^

## Discussion

In this study, we could discern the influence of pyrophosphorylation and phosphorylation on NAGK activity, which was facilitated by the ability to obtain stoichiometrically and site-specifically modified protein. We first expressed pS76-NAGK, using amber codon suppression. Application of chemoselective P-imidazolide chemistry led to the successful generation of ppS76-NAGK in very good yield. In principle, this approach should be readily extendable to other proteins of interest, in which these different phosphorylation modi have been observed. However, the current method to generate pyrophosphoprotein requires the use of organic solvent (DMA) to ensure full conversion and precise protein modification. While refolding of ppS76-NAGK proceeded smoothly, refolding conditions likely have to be adjusted and optimized for other proteins. It would therefore be desirable to develop P-imidazolide reagents that can operate in aqueous environments to preserve protein structure during the course of the reaction.^53^

When pS76-NAGK and ppS76-NAGK were assayed for their kinase activity, a stepwise decrease was observed, compared to wt-NAGK: phosphorylation led to a significant reduction in activity, while pyrophosphorylation resulted in almost complete deactivation. The available crystal structure of wt-NAGK illustrates the close proximity between the ATP-binding site and S76.^16^ Phosphorylation at this site would lead to significant electrostatic repulsion between the modification and the triphosphate moiety of ATP (Figure 7a). This repulsion would decrease ATP-binding and thereby slow down the transfer of the γ-phosphoryl group onto GlcNAc (or MDP). Consistent with this hypothesis, the effect of pyrophosphorylation should be even more pronounced, as was indeed observed.

**Figure 7.**
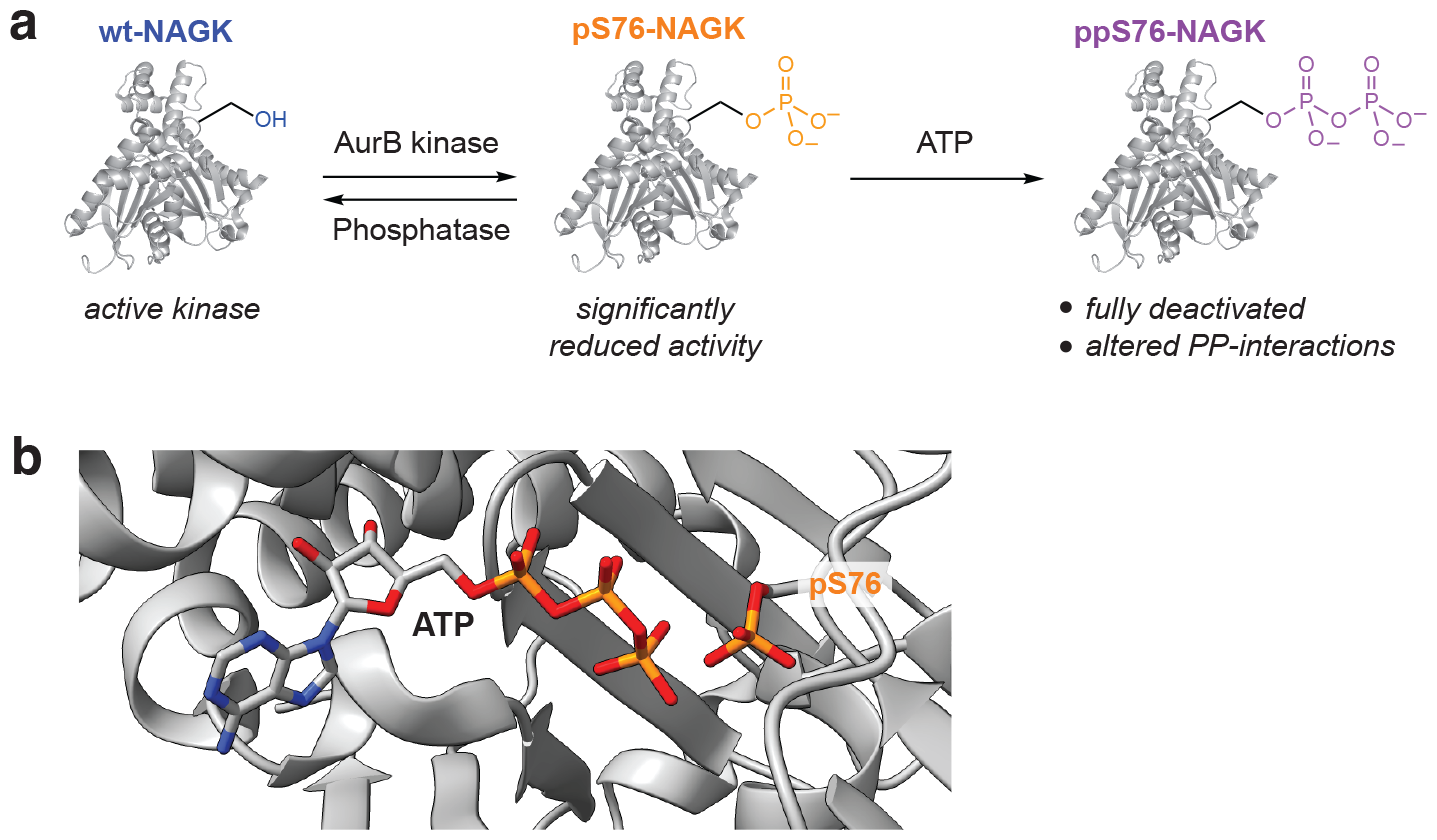
a) Summary of the impact of phosphorylation and pyrophosphorylation on the function of NAGK. b) Structural model of ATP bound to p76-NAGK, based on the NAGK-ADP structure (PDB: 2CH6).

The proximity of the ATP-binding site to pS76 also helps to understand the autocatalytic mechanism, which is proposed for the installation of the pyrophosphate group. While we initially suspected that PP-InsPs would serve as phosphoryl donors for the generation of ppS76-NAGK, these molecules proved to be unreactive. Instead, ATP was the only high-energy phosphoryl donor that promoted the formation of ppS76-NAGK, and the conversion was proportional to ATP concentration. Inspection of the crystal structure of wt-NAGK, into which we modeled a phosphoserine side chain at S76, illustrates how the γ-phosphoryl group of ATP and pS76 are within 3.4 Å of each other, suggesting that an intramolecular phosphoryl-transfer is well possible (Figure 7b).^54^

The initial phosphorylation on serine 76 can be catalyzed by AurB. AurB serves as a central protein kinase across diverse phases of the cell cycle, exerting regulatory control over mitosis and cytokinesis by phosphorylating proteins linked to key structures such as the mitotic spindle, kinetochore, and central spindle.^55—56^

Pyrophosphorylation thereafter is facilitated by the presence of ATP. Given the considerable variability in cellular ATP concentrations, we postulate that this sequential process holds potential for efficiently suppressing the GlcNAc salvage pathway during instances of high local ATP concentrations. Such a mechanism appears well-suited to the nutrient-rich conditions characteristic of such an environment, where reliance on the GlcNAc salvage pathway may be dispensable.^19^

Once the pyrophosphate group is attached to NAGK, it is quite stable towards chemical and biochemical hydrolysis. Consistent with previous observations on protein pyrophosphorylation, extended incubation with active cell lysates did not lead to dephosphorylation of ppS76-NAGK. The more “energetic” pyrophosphoprotein turns out to be much longer lived than its monophosphorylated counterpart. This resilience towards dephosphorylation extends the deactivation period of kinase activity and suggests that an alternate functional outcome may be associated with pyrophosphorylation.

Although limited information is available regarding kinase-independent functions of NAGK, a few recent studies have demonstrated that NAGK can indeed serve as a scaffold by engaging in various protein-protein interactions.^22—25^ The protein-binding partners of catalytically inactive ppS76-NAGK that we identified differed starkly from the binding-partners of the wt-protein. None of the modification-specific interactors had been identified previously, presumably because pyrophosphorylated NAGK only comprises a small fraction of the total protein amount. Notably, ppS76-NAGK associated with several proteins involved in the TCA cycle and mRNA processing. If ppS76-NAGK indeed localizes to mitochondria and has moonlighting functions to regulate cellular ATP production, is something to be investigated in the future.

This resilience of ppS76-NAGK towards dephosphorylation of course also raises the question under which conditions this modification is reversible.^57^ While the lysate conditions cannot capture the complex environment of a cell, the lysate conditions were sufficient to dephosphorylated pS76-NAGK. However, it is possible that the dephosphorylation of ppS76-NAGK requires a specific phosphatase activity within a certain compartment, such as mitochondria or the nucleus. Alternatively, the removal of ppS76-NAGK may depend on a ubiquitin-mediated degradation pathway. A recent study highlighted that pyrophosphorylation of MYC induces polyubiquitination by an E3 ligase, regulating cell survival through a novel “pyrophosphodegron”.^46^ This intriguing mechanism might be relevant to the turnover of ppS76-NAGK as well, as we could identify two E3 ligases that associated with the pyrophosphoprotein.

## Conclusion

In conclusion, this study expands our current understanding of protein pyrophosphorylation. While the change of NAGK structure for the three variants – serine, phosphoserine, pyrophosphoserine – is likely small, the properties have been altered significantly. This granular analysis was only made possible by the synthetic access to stoichiometrically modified NAGK. Notably, several other kinases, including Clk1, GSK3α/β, and NME1/2, have also recently been reported to carry a pyrophosphoserine or pyrophosphothreonine residue in close proximity to the ATP-binding site. It will therefore be of great interest if some of the observations made here about protein pyrophosphorylation – the decrease of catalytic activity, ATP-dependent autopyrophosphorylation, resistance to dephosphorylation – can be extended to these other candidates. We believe that the reported synthetic strategy, using a combination of amber codon suppression and P-imidazolide chemistry, together with the here established analytical tools, will greatly facilitate these investigations in the future.

## Data availability

Methods and Materials as well as supporting figures and tables are available in the Supporting Information.

The mass spectrometry proteomics data have been deposited to the ProteomeXchange Consortium via the PRIDE^1^ partner repository with the dataset identifier PXD049339.

## Acknowledgments

A.C. gratefully acknowledges funding from the DFG (Deutsche Forschunsgemeinschaft) under project number 469186007.

The authors thank Kathryn E. Wellen (University of Pennsylvania) for supplying the human PDA cell lines PANC-1. The authors are grateful to Kathrin Motzny and Lena von Oertzen for their support in molecular cloning and cell culture. Additionally, we gratefully acknowledge the discussions with Christian Stieger and Max Ruwolt on mass spectrometry data. Finally, we than all members of the Fiedler group for the insightful discussions and valuable input.

