## Supporting Information for "An uncommon phosphorylation mode regulates the activity and protein-interactions of N-acetylglucosamine kinase"

^1^: Leibniz-Forschungsinstitut für Molekulare Pharmakologie, Robert-Rössle-Straße 10, 13125 Berlin, Germany

^2^: Institut für Chemie, Humboldt-Universität zu Berlin, Brook-Taylor-Str. 2, 12489 Berlin, Germany

* Corresponding author

This PDF file includes

Supporting Figures S1 to S8

Supporting Methods and references

Table of Contents

1. Supporting Figures……………………….………………………………………………………..1-6
2. General Information…………………………..………...………………………………….6-7
3. Cloning, site directed mutagenesis, expression and purification of recombinant human NAGK……………………………………………………………………..………………..8-11
4. General protocol for protein pyrophosphorylation……………………………………….11
5. General Protocol for NAGK biochemical assays……………………….………………..12
6. General information for the interactome analysis……………………………………13-14
7. Chemical Synthesis and characterization……………………….…………………….....15
8. Q-TOF-MS spectra……………………………………………......…………………....15-17
9. References…………………………………………………………...…………………...…18

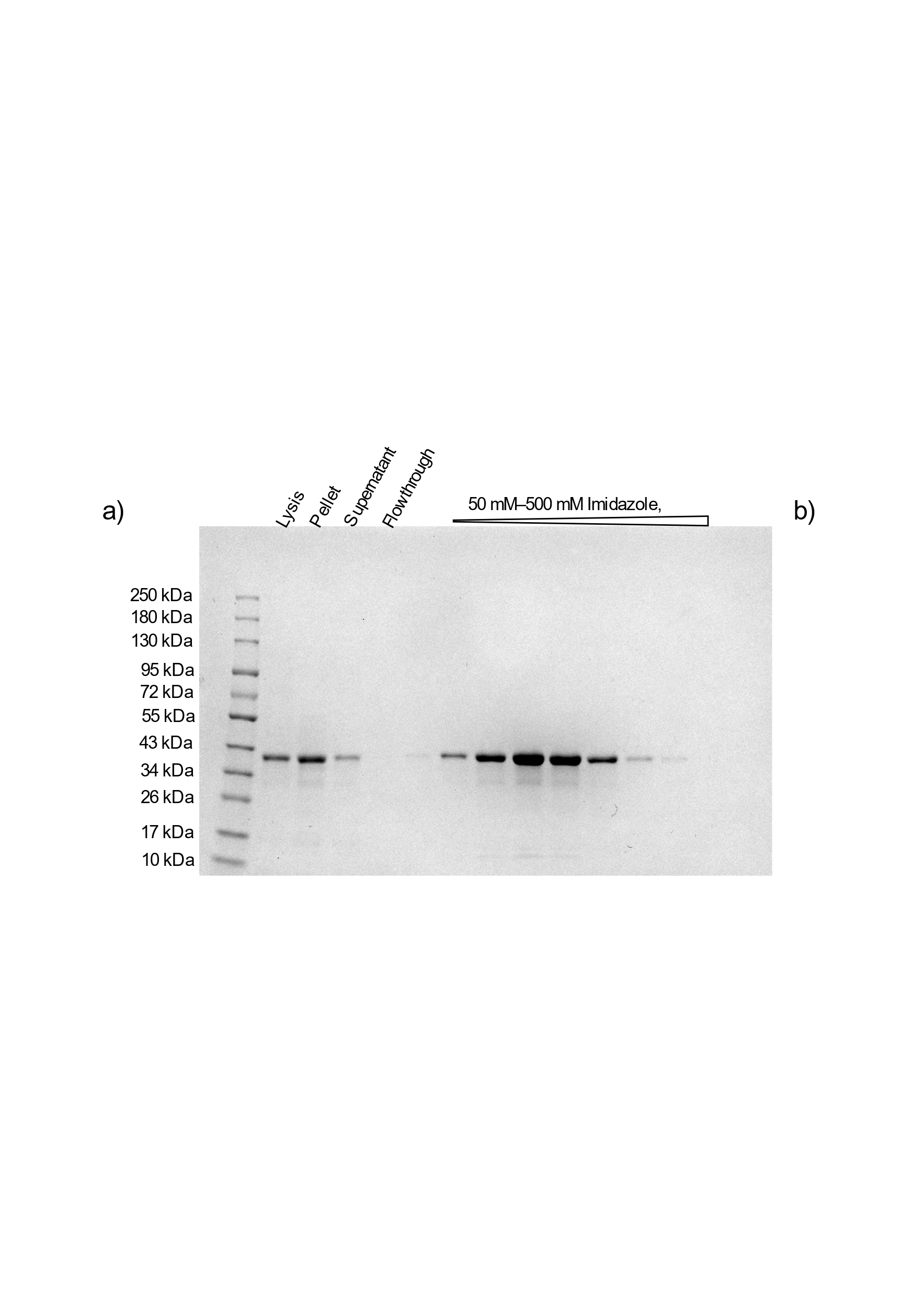

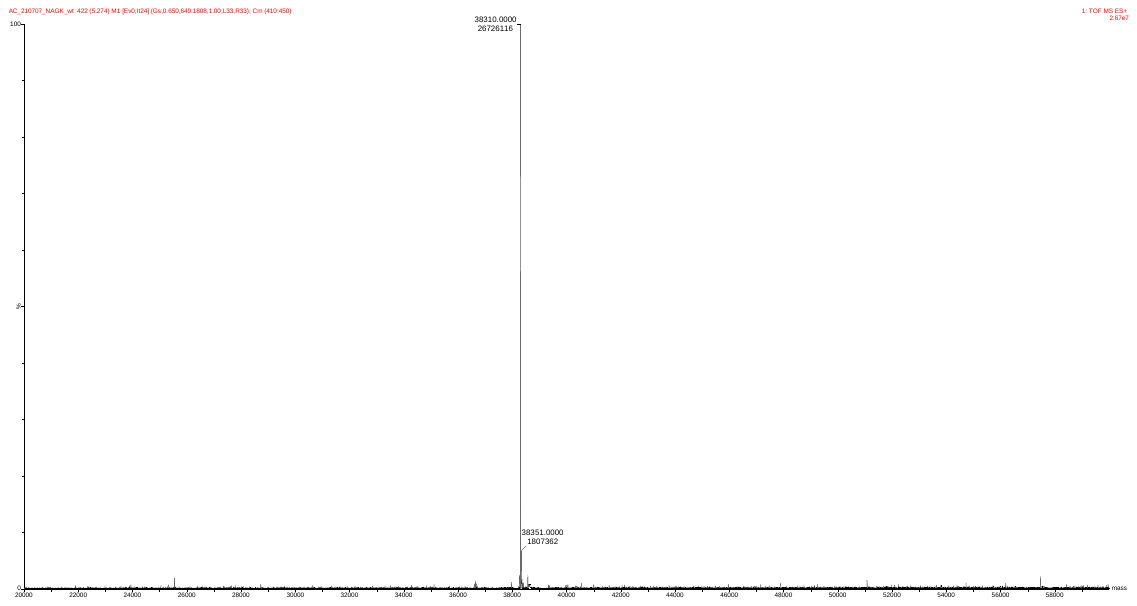
**1. Supporting Figures**

b)

a)

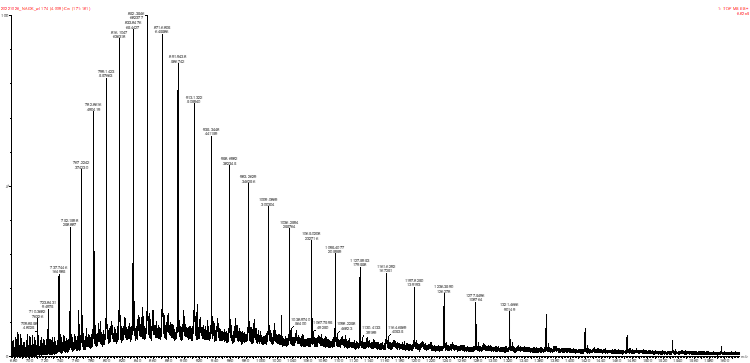

wt-NAGK

38310 Da

**Figure S1.** Recombinant expression of wt-NAGK. a) SDS-PAGE. b) Q-TOF-MS measurement.^1^

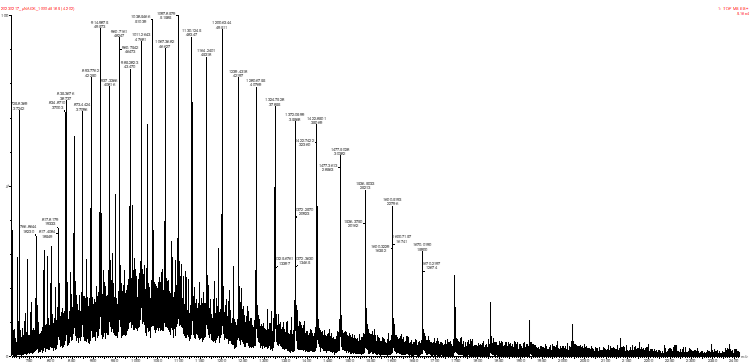

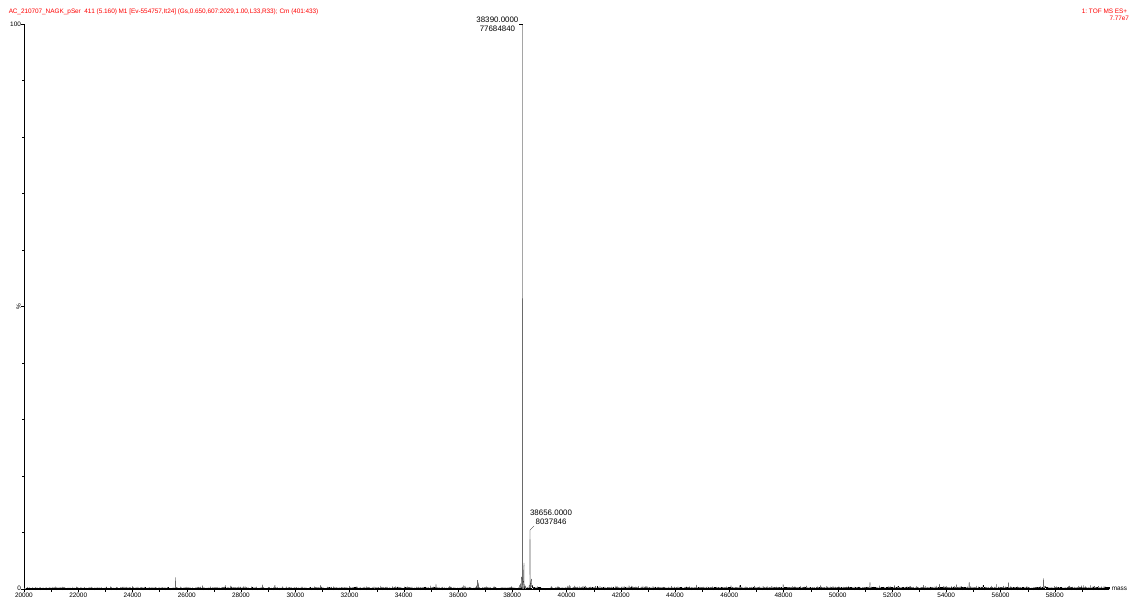

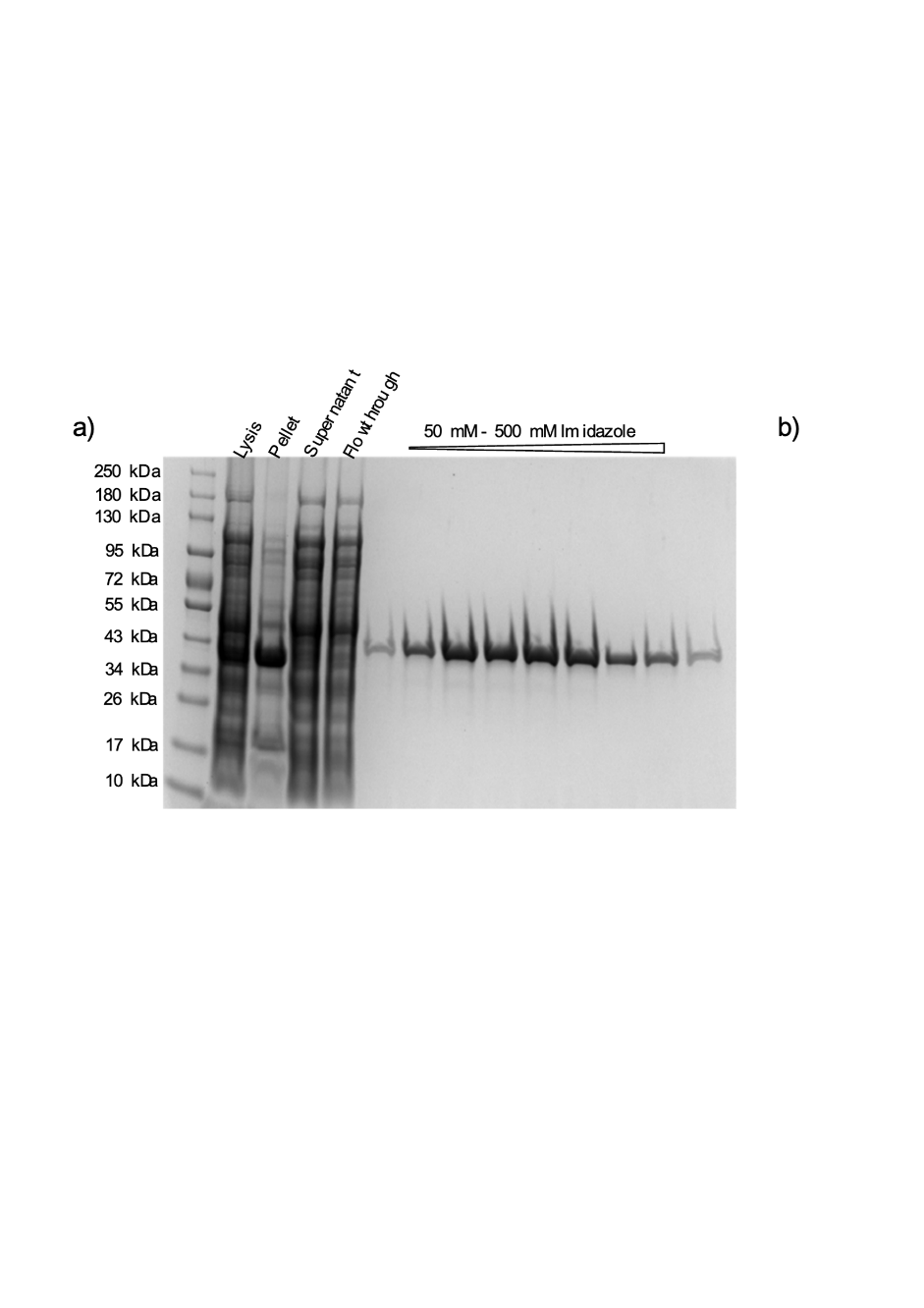

pS76-NAGK

38390 Da

b)

a)

**Figure S2.** Figure S1. Recombinant expression of pS76-NAGK. a) SDS-PAGE b) Q-TOF-MS measurement.^2^

a)

b)

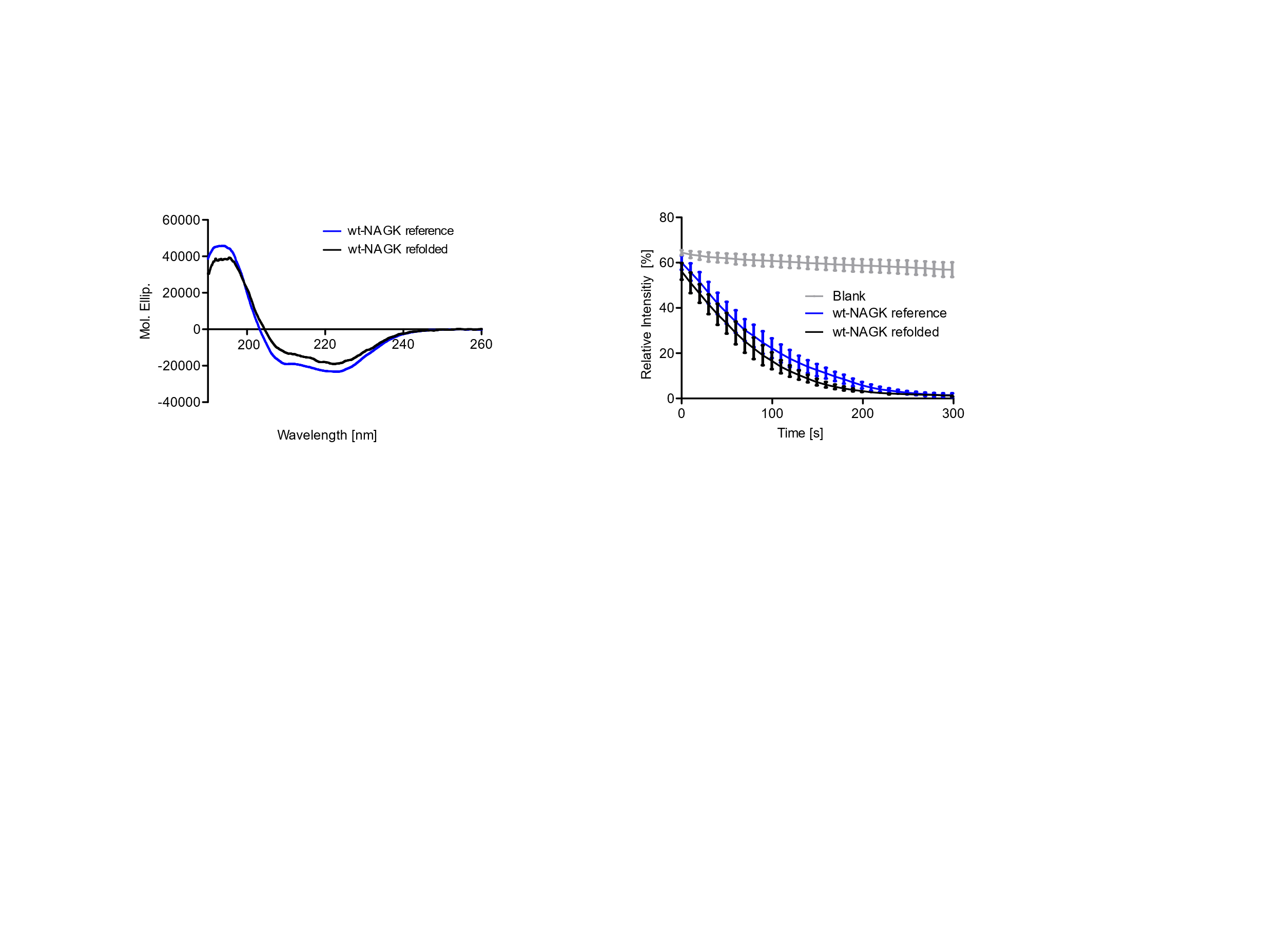

**Figure S3.** Assessing the recovery of wt-NAGK after and before refolding. a) CD-Spectroscopy of wt-NAGK before (in blue) and after refolding (in black). b) GlcNAc activity assay before and after refolding monitored by NADH readout assay.

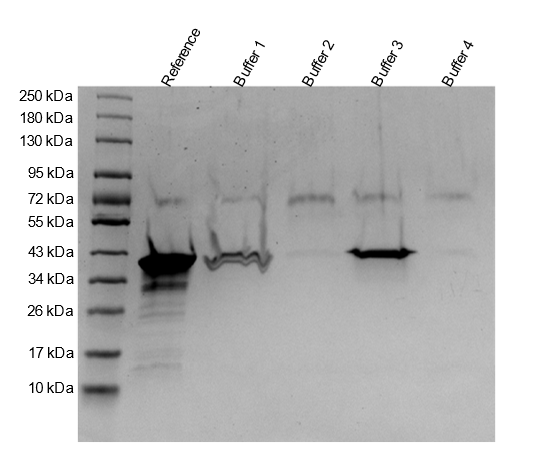

**Figure S4.** Refolding of wt-NAGK analyzed by SDS-PAGE. a) SDS PAGE after applying the refolding protocol on wt-NAGK. b) List of applied refolding buffers. **Buffer 1**: 50 mM Tris-HCl (pH 8.0), 250 mM NaCl, 20 mM DTT, 10 mM EDTA, 0.2% CHAPS, 1 mM GlcNAc. **Buffer 2**: 50 mM Tris-HCl (pH 8.0), 250 mM NaCl, 0.3 mM GSSG, 3 mM GSH, 10 mM EDTA, 0.2% CHAPS, 0.4 M Sucrose, 1 mM GlcNAc. **Buffer 3**: 50 mM Tris-HCl (pH 8.0), 250 mM NaCl, 0.3 mM GSSG, 3 mM GSH, 10 mM EDTA, 0.2% CHAPS, 0.1 M Arginine*HCl, 1 mM GlcNAc. **Buffer 4**: 50 mM Tris-HCl (pH 8.0), 250 mM NaCl, 20 mM DTT, 10 mM EDTA, 0.2% CHAPS, 5 mM GlcNAc, 10% Glycerol.^3,4^

a)

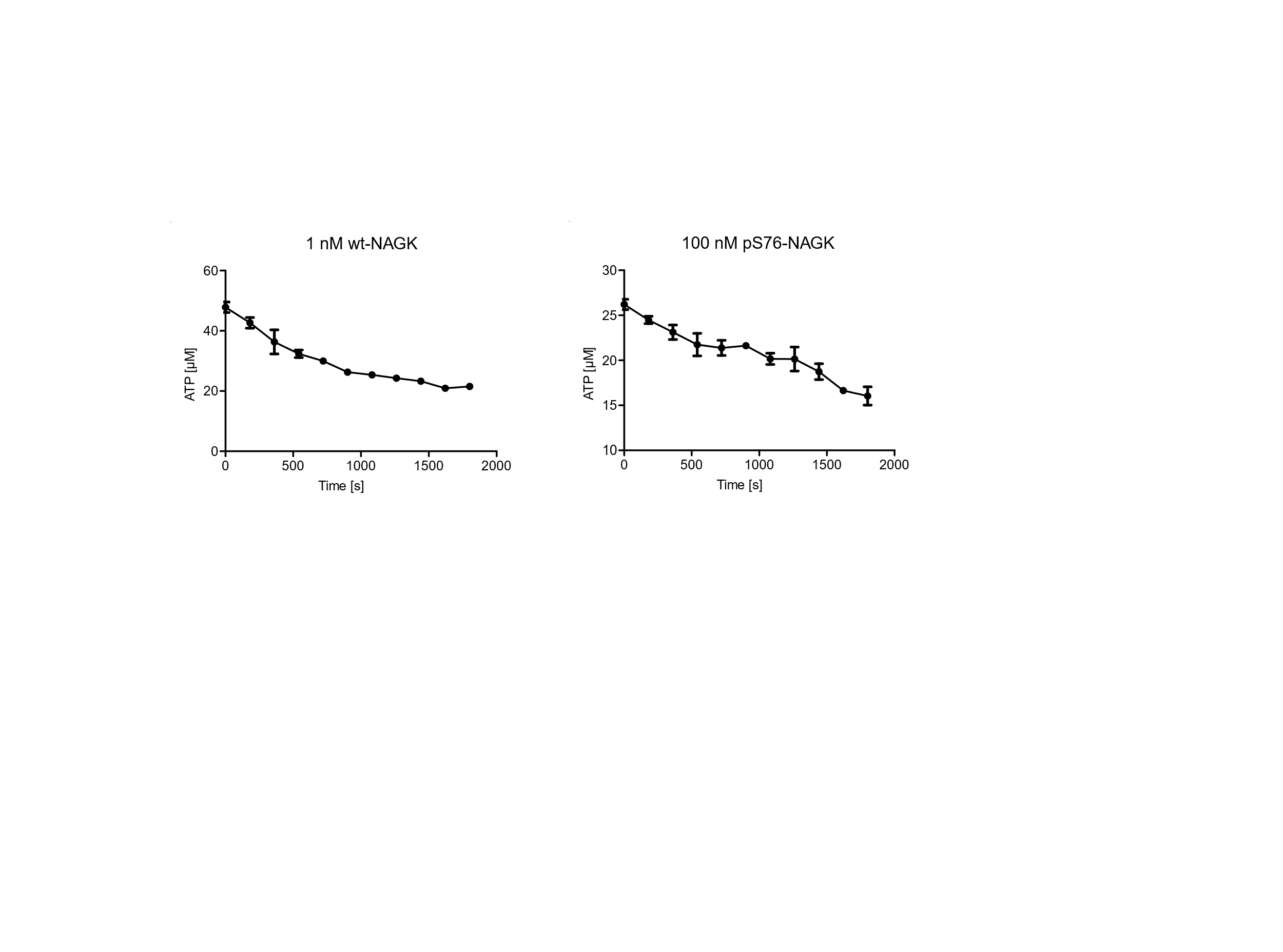

b)

**Figure S5.** Monitoring the time course of wt-NAGK and pS76-NAGK to determine the nmol/min/ng value. a) 1 nM wt-NAGK. b) 100 nM pS76-NAGK.

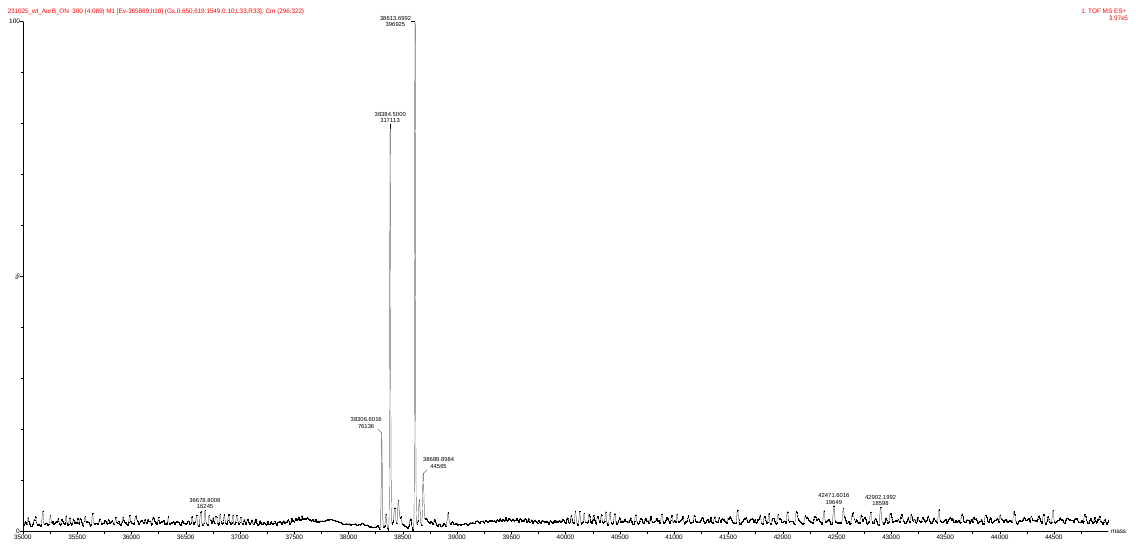

a)

pS76-NAGK

38384 Da

wt-NAGK

38306 Da

b)

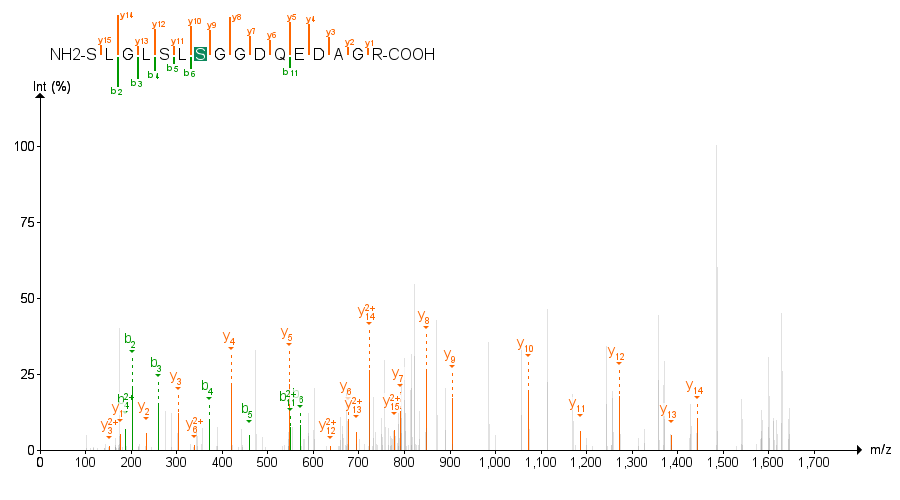

**Figure S5.** MS/MS analysis of AurB treated wt-NAGK localizing phosphorylation on position S76.^5^

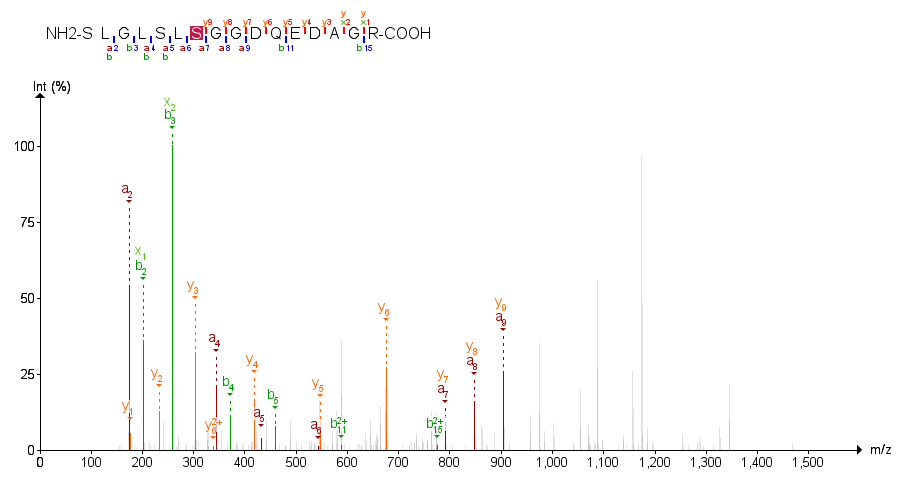

**Figure S6.** MS/MS analysis of ATP treated pS76-NAGK localizing pyrophosphorylation on position S76.^5^

c)

b)

a)

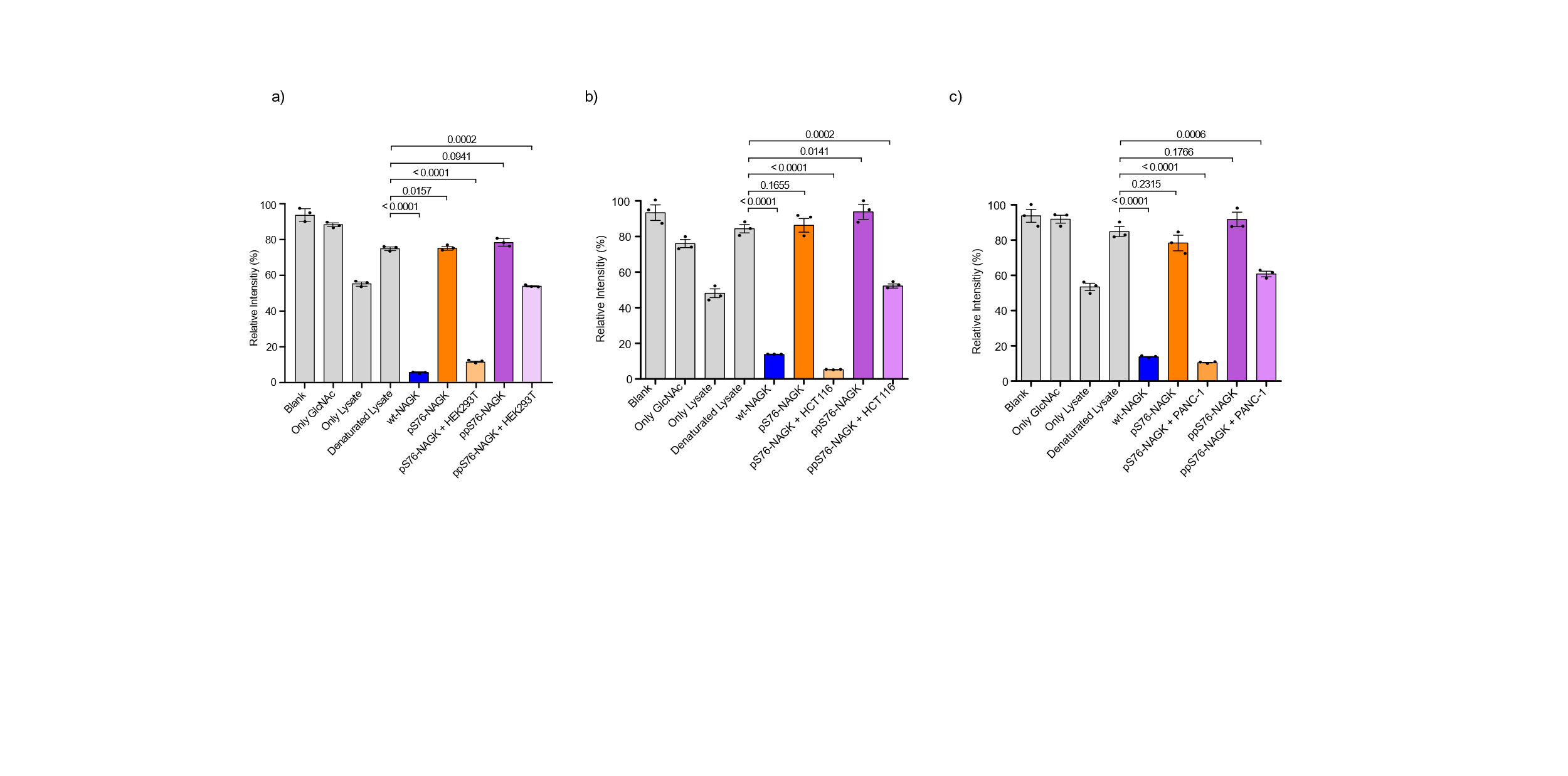

**Figure S7.** Biochemical validation of NAGK (wt, pS76, ppS76) after treatment with a) HEK293T lysates b) HCT116 lysates and b) PANC-1 treatment and subsequent GlcNAc kinase activity assay showed regained activity for pS76-NAGK but not ppS76-NAGK indicating no pyrophosphatase activity in both cell lines. Data presented as mean ±SEM of three technical replicates (N=3). P-values were determined by unpaired t-test analysis.

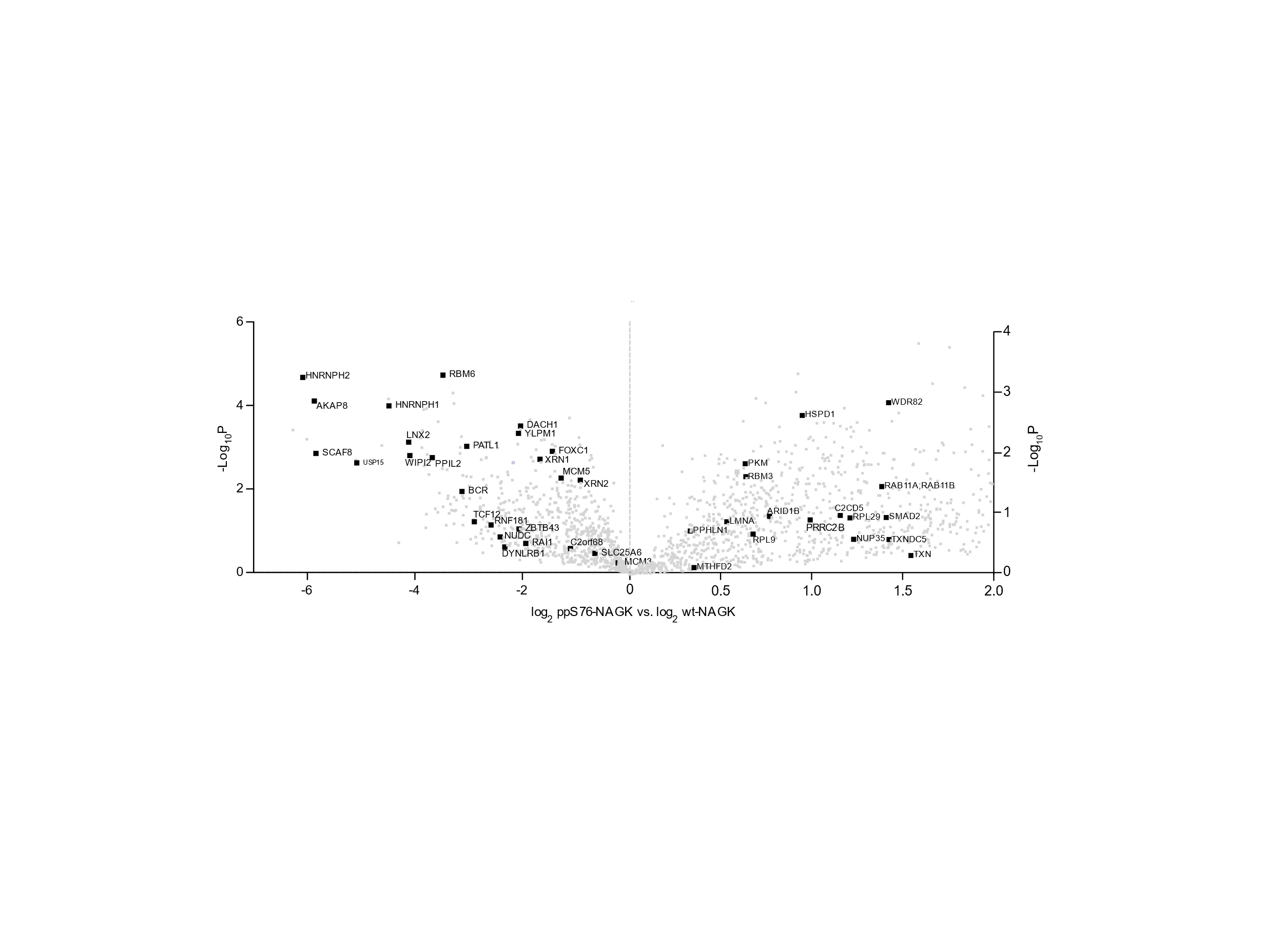

**Figure S8.** Volcano plot depicting LFQ values of ppS76-NAGK versus wt-NAGK after a t-test. The x-axis displays the difference of LFQ values on a log_2_ scale and the y axis shows the -log_10_P value. Left side of the plot represents known interactor (BioGRID database) preferentially enriched with wt-NAGK. Right side of the plot represents known interactors (BioGRID database) preferentially enrichment with ppS76-NAGK.^6,7,8^

**2. General Information**

**2.1 Cell lines**

HEK293T (female human origin), HCT116 (male human origin), and PANC-1 (pancreatic carcinoma-1) cell lines were used in this study. HEK293T and HCT116 cell lines were obtained from ATCC (American Type Culture Collection). PANC-1 cell lines were kindly provided by Kathryn E. Wellen. All cells were tested for mycoplasma prior to use. Cells were grown in 15-cm dishes to 70-80% confluency in Dulbecco’s Modified Eagle’s Medium (DMEM), complemented with 10% FBS, Penicillin-Streptomycin (100 U/mL), Glutamine (2 mM) in a 5% humidified CO_2_ incubator at 37 °C.

**2.2 Chemicals and Solvents**

Commercially available chemicals were purchased from Sigma-Aldrich, Alfa Aesar, Acros Organics, Strem Chemicals, TCI America, Anaspec, Carl Roth GmbH., and Iris. Dichloromethane and THF were dried by passing through an activated alumina column, and acetonitrile, DMSO, and DMF were dried by passing through a column of activated molecular sieves using a Pure Process Technology drying system.

**2.3 Preparative HPLC**

Preparative high-performance liquid chromatography (HPLC) was performed on a Varian system with SD-1 prep solvent delivery system, a ProStar 325 UV-Vis detector and a 440-LC fraction collector, using a Waters XBridge^TM^ 5 µm C18 column (19 × 150 mm).

**2.4 Q-TOF-MS**

High-resolution ESI-MS spectra were recorded on two different instruments: 1) Agilent 6220 TOF Accurate Mass coupled to an Agilent 1200 LC (Agilent Technologies, USA) and were measured at 35 °C between 100–2000 m/z. The used column was an Accucore RP-MS (30 x 2.1 mm; 2.6 μm particle size) eluted with a flow of 0.8 mL/min and the following gradient (A =  H_2_O + 0.1% TFA, B = MeCN + 0.1% TFA), gradient: 5% B 0–0.2 min, 5–99% B 0.2–1.1 min, 99% B 1.1–2.5 min. 2) Agilent Technologies 6230 Accurate Mass TOF LC/MS linked to Agilent Technologies HPLC 1260 Series; Column: Thermo Accucore RP-MS; Particle Size: 2.6 µM Dimension: 30 x 2.1 mm. The following gradient was used: A = H_2_O + 0.1 % formic acid, B = MeCN + 0.1 % formic acid, 5% B 0.0–0.2 min, 5-99% B 0.2–1.1 min, 99% B 1.1–3.6 min, 5% B 3.6–4.9 min. Flow rate: 0.8 mL/min; UV-detection: 220 nm, 254 nm, 300 nm.

**2.5 Intact protein MS**

Intact proteins were analyzed using a Waters H-class instrument equipped with a quaternary solvent manager, a Waters sample manager-FTN, a Waters PDA detector and a Waters column manager with an Acquity UPLC protein BEH C4 column (300 Å, 1.7 μm, 2.1 mm x 50 mm). Proteins were eluted at a column temperature of 80 °C with a flow rate of 0.3 mL/min. The following gradient was used: A = H_2_O + 0.01% formic acid, B = MeCN + 0.01% formic acid. 5–95% B 0–6 min at 40 °C. Mass analysis was conducted with a Waters XEVO G2-XS QTof analyzer. Proteins were ionized in positive ion mode applying a cone voltage of 40 kV. Raw data was deconvoluted with MaxEnt.

**3. Cloning, site directed mutagenesis, expression and purification of recombinant human NAGK**

**3.1 Cloning**

A gene sequence encoding for human NAGK (full-length, Uniprot Q9UJ70) was purchased using Thermo Fisher’s GeneArt service. The sequence was codon optimized for expression in *E. coli* and contains a NdeI (at initial ATG) and XhoI (after the stop codon) restriction site. The MINPP1 gene was cloned into the vector pET-21a using the NdeI and XhoI restriction sites. The resulting plasmid (pET-15b-NAGK) encodes an N-terminal His-tag followed by NAGK. For plasmid preparation the E. coli Top10 strain was used.

**3.2 Site directed mutagenesis**

Site directed mutagenesis was performed on a Bio-Rad C1000 Touch Thermal Cycler machine. Plasmid DNA harboring the corresponding NAGK ORF was extracted from an overnight culture of the pET21-NAGK vector in TB-Amp using a QIAGEN Miniprep kit. Single point mutations were installed by employing the Forward (5’ – GCTGGGTGAGTAGAATCCTGCTG) and reverse (5’ – ATCACGCGACCTGTCTTT) primer. 50 µl PCR reactions were performed on a Bio-Rad C1000 Touch Thermal Cycler following the NEB Phusion High-Fidelity DNA polymerase protocol using the following temperature program: 5 min 98 °C → 30 s 98 °C → 3 min 62 °C → Cycle to step 2 30× → 5 min 72 °C.

**Mutagenesis Primers**

Forward: AGCCTGTAGGGTGGTGATCAAGAAG

Reverse: ACCACCCTACAGGCTCAGACC

**Sequencing Primer**

T7 Forward: 5’ – TAATACGACTCACTATAG – 3’

**3.3 Plasmid Transformation**

**Heat-shock procedure**

Roughly 100 ng DNA were added to competent *E. Coli* cells and incubated on ice for 30 min. After the heat shock (40 s at 42 °C), the cells were kept on ice for 5 min. Subsequently, 500 μL SOC outgrowth medium was added and the cells incubated for 30 min at 37 °C before being plated onto LB plates supplemented with the required antibiotic, and grown overnight at 37 °C. Colonies were picked and inoculated for overnight cultures in LB supplemented with the required antibiotic (Ampicillin 50 μg/mL, Chloramphenicol: 25 μg/mL) out of which glycerol stocks were prepared. The obtained plasmids were extracted with QIAprep 2.0 Spin Miniprep Kit following manufacturer’s procedure and sent in for sequencing.

**Electroporation procedure**

Electrocompetent BL21 (DE3) ∆serB cells (25 μL) were combined with 10.0 ng of DNA plasmids in electroporation cuvettes. Pulsing was performed using a Gene Pulser X-cell system (BioRad) with default *E. coli* settings (Voltage: 1800 V, Capacitance: 25 μF, Resistance: 200 Ω, Gap length: 1.0 mm). Post-pulsing, pre-warmed SOC medium was added within 30 s. Electroporated cell suspensions were transferred to 5 mL tubes, incubated (37 ˚C, 200 rpm) for 1 h, and plated onto LB plates with appropriate antibiotics.

**Gel Electrophoresis and Staining**

Protein samples were combined with 4x Laemmli sample buffer (Bio-Rad) containing 10% β-mercaptoethanol and heated to 95 °C for 8 min. Following heating, the samples were cooled and consolidated by centrifugation at 5000 x g for 1 minute before loading onto Bio-Rad Mini-Protean TGX Stain-Free precast 4–20% gradient gels with 10x 30 µL wells. Gel electrophoresis was conducted using a Bio-Rad PowerPac HC 300 W power source set to 150 V for 1 h or until the loading dye migrated off the gel. Subsequently, gels underwent three 5-min washes with deionized water (D.I. H_2_O) and were stained for 1 h with GelCode^TM^ Blue colloidal Coomassie stain G-250 from ThermoFisher Scientific. Finally, gels were de-stained through multiple washes with D.I. water.

**3.4 Protein Expression**

**Expression of wildtype-NAGK**

*E. Coli* BL21 (DE3) harboring the required pET21a NAGK vector were inoculated in 10 mL LB-Amp overnight at 37 °C. The overnight culture was diluted to a final OD_600_ of 0.05 and grown to OD_600_ of 0.7 at 37 °C. The temperature was switched to 18 °C and expression induced with 1 mM isopropyl β-D-1-thiogalactopyranoside (IPTG). After 20 h expression, the cells were pelleted by centrifugation (3000 x *g*, 10 min, 4 °C), washed with LB medium and centrifuged again. The cell pellet was stored at -80 °C after removal of the supernatant water by another centrifugation.

**Expression of pS76-NAGK *via* Amber Codon Suppression**

*E. Coli* BL21 (DE3) ∆serB harboring the required pET21a (Amp^R^) NAGK and pKW1-Sep (Camp^R^) vector were inoculated in LB-Amp-Camp overnight at 37 °C. The overnight culture was diluted to a final OD_600_ of 0.05 and grown to OD_600_ of 0.7 at 37 °C in the presence of Amp and Camp. The temperature was switched to 18 °C and expression induced with 1 mM isopropyl β-D-1-thiogalactopyranoside (IPTG) and 2 mM (L)-O-phosphoserine (pH 7.0). After 20 h expression, the cells were pelleted by centrifugation (3000 x *g*, 10 min, 4 °C), washed with LB medium and centrifuged again. The cell pellet was stored at -80 °C after removal of the supernatant water by another centrifugation.

**General protein purification procedure**

The frozen cell pellet was thawed and resuspended in 8 mL lysis buffer (50 mM Tris-HCl (pH 7.8), 150 mM NaCl, 50 mM Imidazole) per 1 g of wet weight and supplemented with lysozyme, DNase I and 1 tablet of complete protease inhibitor (purchased from Sigma-Aldrich). After 30 min of incubation on ice the cell extract was lysed with a microfluidizer^TM^ LM10 at 15000 psi with five iterations. The cell debris was removed by centrifugation (30000 x g, 30 min, 4 °C) and the supernatant lysate filtered (VWR vacuum filter, PES, 0.45 μm). Recombinantly expressed protein were purified on a FPLC system (NGC Quest 10 Chromatography System, Bio-Rad). For purification, the lysate was loaded onto a Co-NTA column that was equilibrated with lysis buffer at a flowrate of 2.5 mL/min. The protein was eluted with a 0–100% gradient of elution buffer (50 mM Tris-HCl (pH 7.8), 150 mM NaCl, 500 mM Imidazole). The volume of the fractions that contained the desired protein were reduced by spin filtration through a 10 kDa cut-off filters and dialyzed overnight against dialysis buffer (50 mM Tris-HCl (pH 7.8), 150 mM NaCl, 1 mM DTT, and 10% Glycerol). Protein concentrations were determined using a PierceTM BCA Protein Assay Kit. Recombinantly expressed protein were purified on a FPLC system (NGC Quest 10 Chromatography System, Bio-Rad).

**wt-NAGK**

Strong cation exchange FPLC purification: A pre-packed 1 mL GE Healthcare Hi Trap^TM^ SP HP cation exchange column was equilibrated with 50 mM Tris HCl and 200 mM NaCl in D.I. H_2_O, pH 8.0. A gradient of 0–250 mM NaCl in 50 mM Tris HCl Buffer, pH 8.0 over 30 column volumes were applied. All fractions were combined and concentrated to in a 3 kDa MWCO centrifugation filter 2x (3214 x g, 20 min, 15 °C, 15 mL capacity) without exchanging the buffer. Protein concentration: 33 mg/L.

Sequence:

AAIYGGVEGGGTRSEVLLVSEDGKILAEADGLSTNHWLIGTDKCVERINEMVNRAKRKAGVDPLVPLRSLGLSL**S**GGDQEDAGRILIEELRDRFPYLSESYLITTDAAGSIATATPDGGVVLISGTGSNCRLINPDGSESGCGGWGHMMGDEGSAYWIAHQAVKIVFDSIDNLEAAPHDIGYVKQAMFHYFQVPDRLGILTHLYRDFDKCRFAGFCRKIAEGAQQGDPLSRYIFRKAGEMLGRHIVAVLPEIDPVLFQGKIGLPILCVGSVWKSWELLKEGFLLALTQGREIQAQNFFSSFTLMKLRHSSALGGASLGARHIGHLLPMDYSANAIAFYSYTFSLEHHHHHH

**pS76-NAGK**

Strong cation exchange FPLC purification: A pre-packed 1 mL GE Healthcare Hi Trap^TM^ SP HP cation exchange column was equilibrated with 50 mM Tris HCl and 200 mM NaCl in D.I. H_2_O, pH 8.0. A gradient of 0–250 mM NaCl in 50 mM Tris HCl Buffer, pH 8.0 over 30 column volumes were applied. All fractions were combined and concentrated to in a 3 kDa MWCO centrifugation filter 2x (3214 x g, 20 min, 15 °C, 15 mL capacity) without exchanging the buffer. Protein concentration: 7.5 mg/L.

Sequence:

AAIYGGVEGGGTRSEVLLVSEDGKILAEADGLSTNHWLIGTDKCVERINEMVNRAKRKAGVDPLVPLRSLGLSL**pS**GGDQEDAGRILIEELRDRFPYLSESYLITTDAAGSIATATPDGGVVLISGTGSNCRLINPDGSESGCGGWGHMMGDEGSAYWIAHQAVKIVFDSIDNLEAAPHDIGYVKQAMFHYFQVPDRLGILTHLYRDFDKCRFAGFCRKIAEGAQQGDPLSRYIFRKAGEMLGRHIVAVLPEIDPVLFQGKIGLPILCVGSVWKSWELLKEGFLLALTQGREIQAQNFFSSFTLMKLRHSSALGGASLGARHIGHLLPMDYSANAIAFYSYTFSLEHHHHHH

**4. General protocol for protein pyrophosphorylation**^10^

The storage buffer for all protein aliquots used in pyrophosphorylation reactions was removed and exchanged for Milli-Q H_2_O by at least 5 cycles of spin desalting. Amicon® Ultra Centrifugal Filters from Millipore Sigma with capacities of 15 mL or 500 μL were used for concentration and desalting of protein solutions. Steps which call for heating and agitation of protein samples were performed in an Eppendorf^TM^ ThermoMixer®, equipped with a heated lid. Solutions made with DMA were passed through a 0.45 μm PTFE syringe filter before use. Photochemical reactions were conducted using an Atlas Photonics Lumos 43 light source with an optical output of 200 mW/cm^2^.

**4.1 Protocol for protein pyrophosphorylation and refolding**

A solid sample of P-imidazolide reagent was supplemented with DMA containing 340 mM ZnCl_2_ to yield a 68 mM P-imidazolide solution. Subsequently, a Protein low binding 0.5 mL tube containing a 5.0 µL aliquot of 10.0 µg/µL NAGK in Milli-Q H_2_O received 45.0 µL of the freshly prepared P-Imidazolide solution. The resultant clear yellow solution (comprising 61.2 mM P-imidazolide, 26 µM protein, and 306 mM ZnCl_2_, in a 1:9 ratio of H_2_O to DMA) underwent incubation at 45 °C with 1000 rpm shaking for 21 h. A 50 µL aliquot was quenched by dilution into 450 µL of solubilization buffer (50 mM Tris-HCl (pH 8.0), 6.0 M Guanidine chloride. 0.2% CHAPS, 10 mM DTT) and further incubated for 2 h at 37 °C. The solution was treated with refolding buffer (composed of 50 mM Tris-HCl pH= 8.0, 250 mM NaCl, 0.3 mM GSSG, 3 mM GSH, 10 mM EDTA, 0.2% CHAPS, 0.1 M Arginine*HCl, 1 mM GlcNAc) and dialyzed overnight at 4 °C. Subsequently, the samples underwent concentration via multiple rounds of centrifugation in a 30 kDa MWCO centrifuge filter unit with a 15 mL capacity (20 min, 3,214 x *g*, 17 ˚C) to obtain the concentrated protein. Finally, NAGK was subjected to 360 nm light irradiation for 1 h and subsequently analyzed by Q-TOF-MS.

**4.2 Circular Dichrorism (CD) Spectroscopy:**

CD spectra were obtained on a Jasco J-720 spectropolarimeter using Jasco J-700 series control driver software, version 1.08.00 [Build 3]. Data was analyzed with Jasco Spectra Analysis software, version 1.53.04 [Build 1]. Spectra were taken in a Hellma 100-QS Quartz SUPRASIL® Cuvette with a 1.0 mm path length. Samples were prepared by exchanging into Phosphate buffered Fluoride (PBF) buffer (consisting of 154 mM NaF, and 10 mM Na_2_HPO_4_ in Milli-Q H_2_O adjusted to pH 7.4, made with 99.99% NaF (trace metals basis, from Sigma-Aldrich)) by at least five cycles of spin desalting (18,000 x *g,* 20 min, rt) in 500 µL 10 kDa MWCO centrifugal filters, by diluting the sample to 500 µL with PBF between spins. Sample protein concentrations were adjusted to 10 µM as determined by UV absorbance at 280 nm.

**5. General Protocol for NAGK biochemical assays**

**5.1 NAGK kinase assay**

NADH readout assay:

Purified NAGK (10 nM) was added to a reaction mixture containing 50 mM HEPES, 2 mM ATP, 2.2 mM ATP, 0.2 mM β-NADH, 1.1 mM PEP, 10 mM MgCl_2_ 10 units lactic dehydrogenase and 7 units pyruvate kinase to a final volume of 100 µL. The reaction was vortexed immediately and decrease in absorbance of NADH at 340 nm was then recorded for 5 min. UV signals were read out with a TECAN Infinite M Plex plate reader.

Kinase Glo assay:

Purified NAGK (1 nM–1000 nM) was added to a reaction mixture containing 40 µM ATP, 50 mM HEPES, 100 mM NaCl, 10 mM MgCl_2_, 1 mM DTT, and 35 µM GlcNAc. After 1 h at 37 °C, 15 µL of *Promega* Kinase-Glo Plus^®^ reagent were added and the luminescence read out with a *Tecan* Infinite M Plex reader using 100 ms exposure time after 10 min of equilibration. Luminescence signals were read out with a TECAN Infinite M Plex plate reader.

Autopyrophosphorylation assay

pS76-NAGK (10 µM) was incubated in kinase buffer (50 mM HEPES pH= 7.4, 100 mM NaCl, 10 mM MgCl_2_, 200 µM – 2 mM ATP, 1 mM DTT) at 37 ^o^C overnight. Subsequently, the sample was digested by trypsin in solution and analyzed by MS/MS. Relative quantification was determined by the Software FreeStyle^TM^ 1.7 integrating the MS1 intensities relative to the background.

**6. General information for interactome analysis**

**6.1 HEK293T cell culture and lysate processing**

HEK293T cells were grown in 15-cm dishes to 70–80% confluency in Dulbecco’s Modified Eagle’s Medium (DMEM), complemented with 10% FBS, Penicillin-Streptomycin (100 U/mL) and Glutamine (2 mM). Cells were washed twice with ice-cold DPBS (10 mL) and lysed by sonication (IKA Labortechnik, U200S control, 0.5 cycles, 50% intensity, 5 rounds) in 50 mM TBS buffer. supplemented with phosphatase and protease inhibitors (Roche PhosStopTM and cOmpleteTM EDTA-free protease inhibitor cocktail). The cells were scraped off, transferred to protein low binding microcentrifuge tubes and incubated on ice for 10 min. The lysate was then centrifuged at 4°C for 10 min at 17,900 g. The supernatants were combined and lysate protein concentration was determined using Pierce^TM^ Coomassie (Bradford) protein-assay-kit.

**6.2 Affinity Capture Experiments for Proteomic Analysis**

All steps were conducted at 4 °C. A suspension of Nickel-beads (50 µL) were washed three time with 1 mL Milli-Q H_2_O and three time with 1 mL TBS Buffer (50 mM Tris-HCl pH= 7.5, 150 mM NaCl.) supplemented with 2 mM MgCl_2_ and 2 mM MnCl_2_. Subsequently, 25 µg of recombinant NAGK (wildtype, S76pS, S76ppS) in 100 µL TBS Buffer was added to the beads and incubated with constant rotation for 1 h at 4°C. Beads were centrifuged at 2000 x *g* and the supernatant was discarded. The beads were washed three times with 1 mL TBS buffer and 1 mg of HEK293T cell lysate in TBS buffer was added to the beads and incubated for 3 h at 4 °C under rotation. Upon this time, the beads were centrifuged at 2000 x *g*, the supernatant was discarded and then they were washed three times with 1 mL TBS buffer. Lastly, proteins were incubated with elution buffer (50 mM Tris-HCl (pH 7.5), 150 mM NaCl, 500 mM Imidazole) for 1 h at 4 °C under constant rotation. After centrifugation at 2000 x *g*, the supernatant was collected and lyophilized.

**6.3 In solution tryptic digestion**

Lyophilized samples were resolubilized in 100 µl Buffer (50 mM TEAB, 6 M Urea) and diluted to 2 M Urea. Samples were subsequently reduced and alkylated with 5 mM TCEP and 20 mM Iodoacetamide (IAA) at 37 °C for 1 hour. Trypsin was added at an enzyme to protein ration of 1:100 (w/w) to digest overnight at 37 °C. Trypsin was then quenched with 1% FA (endconcentration) and centrifuged for 10 min at 20000 x g. The supernatant was collected and desalted using Sep-PAK C18 cartridges and lyophilized. Peptides were quantified by using BCA quantification.

**6.4 Liquid Chromatography and mass spectrometry**

LC-MS/MS analysis was performed using an UltiMate 3000 RSLC nano LC system coupled on-line to an Orbitrap Fusion or Lumos mass spectrometer (Thermo Fisher Scientific). For sample loading a PepMap C-18 trap-column (Thermo Fischer Scientific) of 0.075 mm ID x 50 mm length, 3 μm particle size and 100 Å pore size was used. The loading mobile phase A contained 1% acetonitrile and 0.05% TFA acid in water, and mobile phase B 0.05% TFA acid in acetonitrile. Reversed-phase separation was performed using a 50 cm analytical column (in-house packed with Poroshell 120 EC-C18, 2.7 μm, Agilent Technologies) with mobile phase A contained 0.1% formic acid in water, and mobile phase B 0.1% formic acid in acetonitrile. The gradient started with 4% buffer B reaching 40% buffer B in 95 min, with total run time of 120 min including column wash and equilibration. MS1 scans were performed in the orbitrap using 120000 resolution; MS2 scans were acquired in the ion trap with an AGC target of 10000 and maximum injection time of 35 ms, charge state 2-4 enable for MS2.

**6.5 Identification, quantification and statistics of proteomics data**

Raw data were analyzed and processed using MaxQuant software version 1.6.2.6a. Analysis was done with standard settings. Search parameters included two missed cleavage sites, fixed cysteine carbamidomethyl modification, and variable modifications including methionine oxidation and N-terminal protein acetylation. The peptide mass tolerance was 4.5 ppm for MS scans and 20 ppm for MS/MS scans. The match between runs option was enabled. Database search was performed using Andromeda against the Human UniProt/Swiss-Prot database (October 2016) with common contaminants. The false discovery rate (FDR) was set to 1% at both the peptide and protein level. Protein quantification was done based on razor and unique peptides. Label-free quantification was enabled. Bioinformatic analysis was carried out using Perseus software version 1.6.7.0. Proteins were filtered to exclude reverse database hits, potential contaminants, and proteins only identified by site. Proteins were further filtered by rows, requiring a valid value for at least two proteins out of four technical replicates. Data was imputed using Perseus default parameters, 0.3 width and 1.8 down shift. Volcano plots were generated using a t test (number of randomizations: 250).

**7. Chemical Synthesis and Characterization**

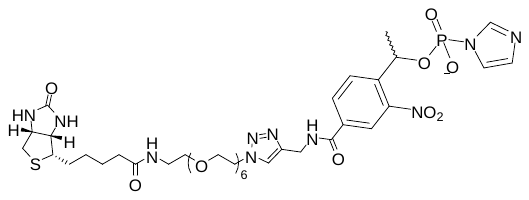

Biotin-PEG_6_-trayole-NPE-(1H-imidazol-1-yl)phosphonate was synthesized as previously described.^9^ Spectral data matches previously reported values.

**8. Q-TOF-MS spectra**

wt-NAGK

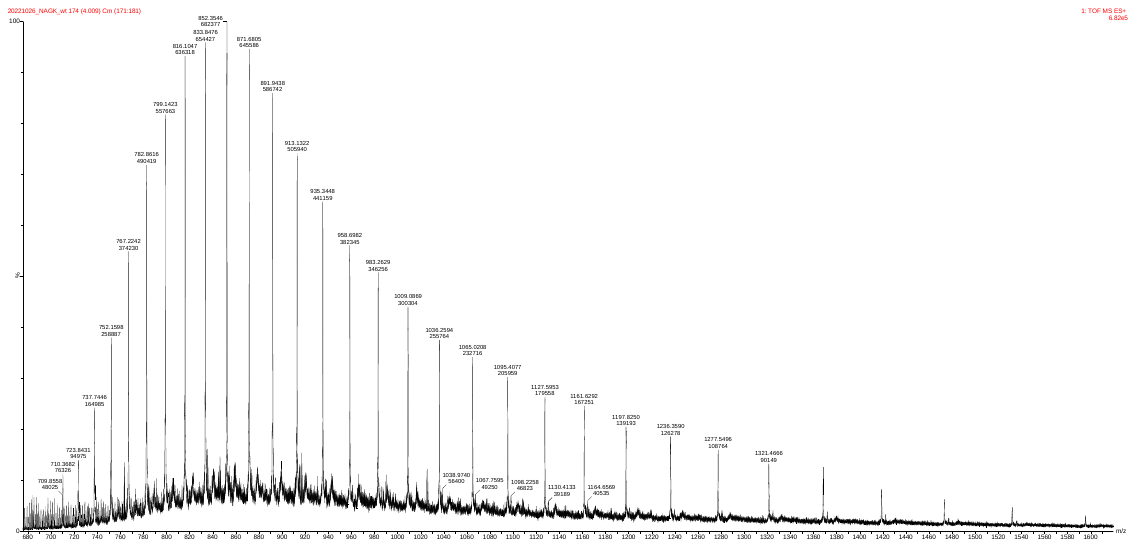

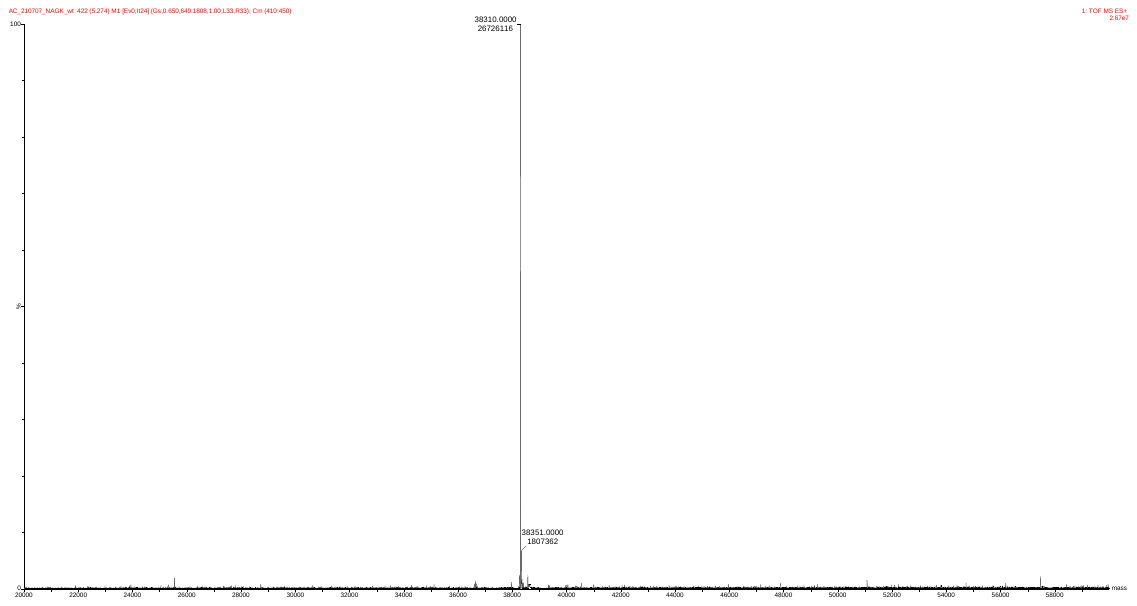

pS76-NAGK

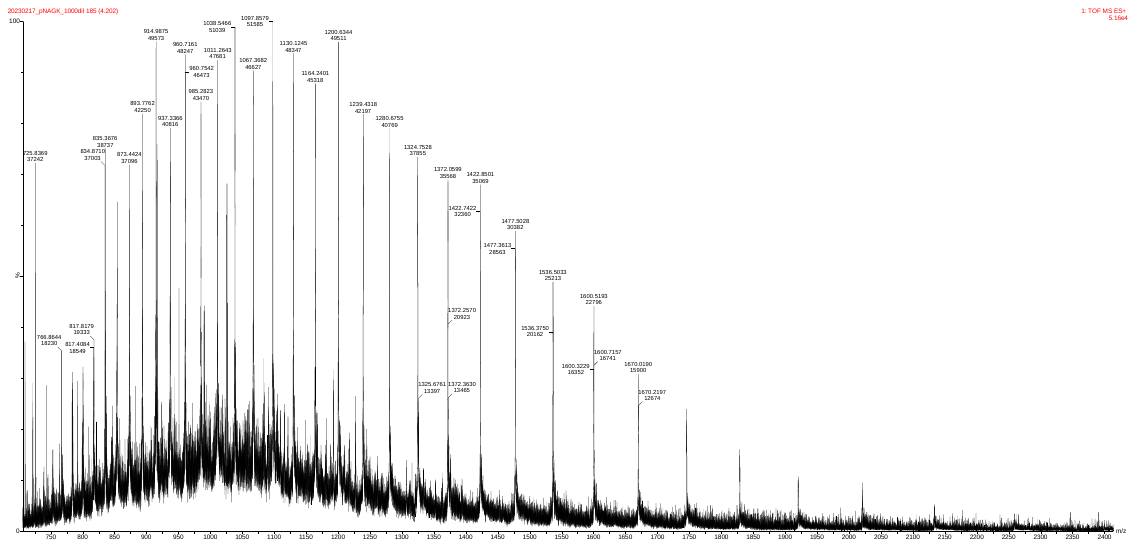

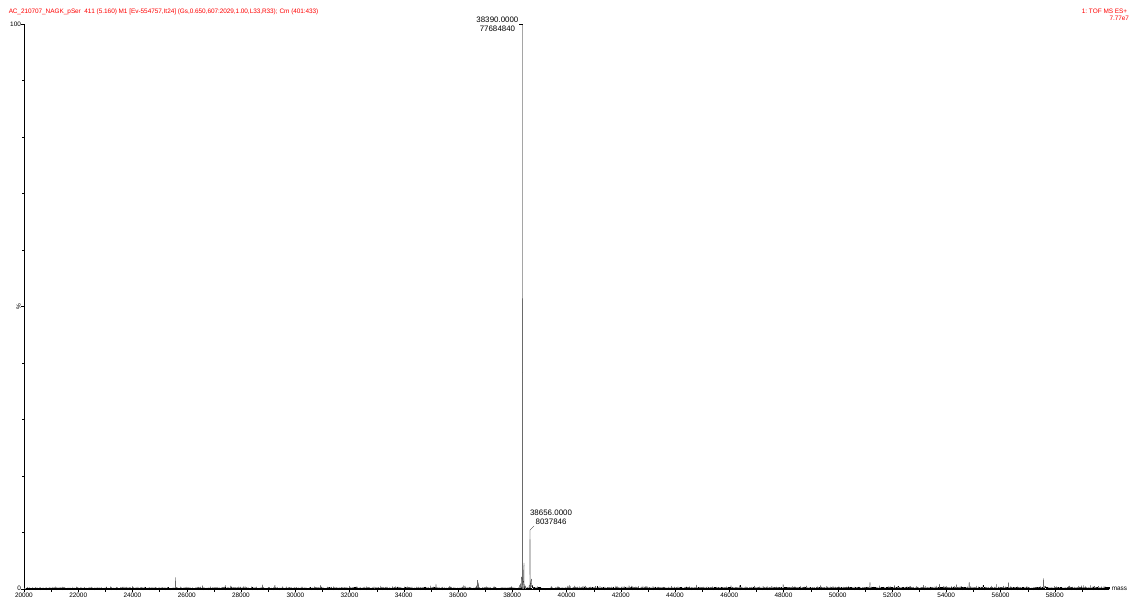

R-ppS76-NAGK

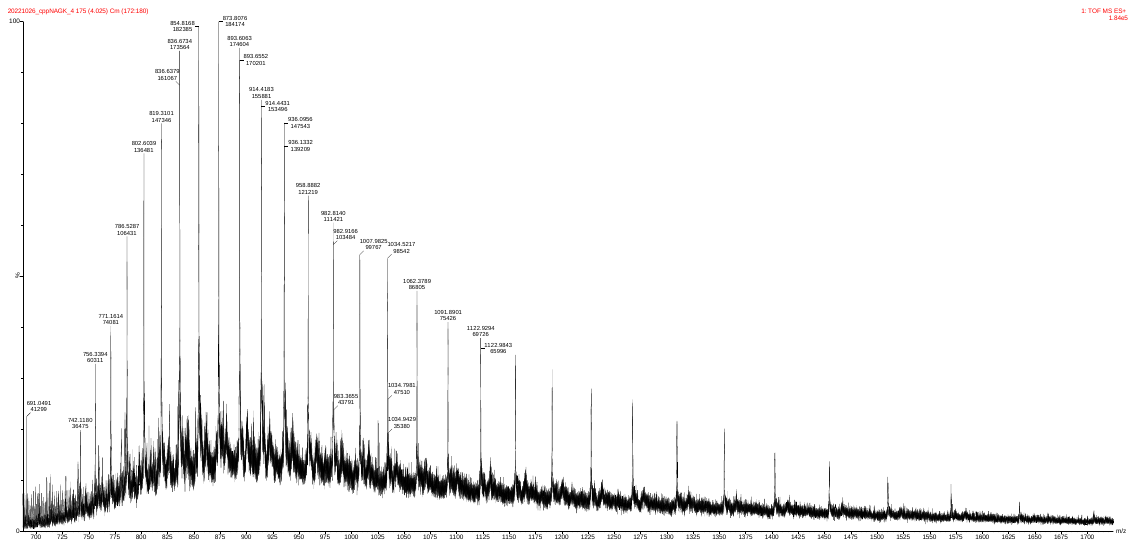

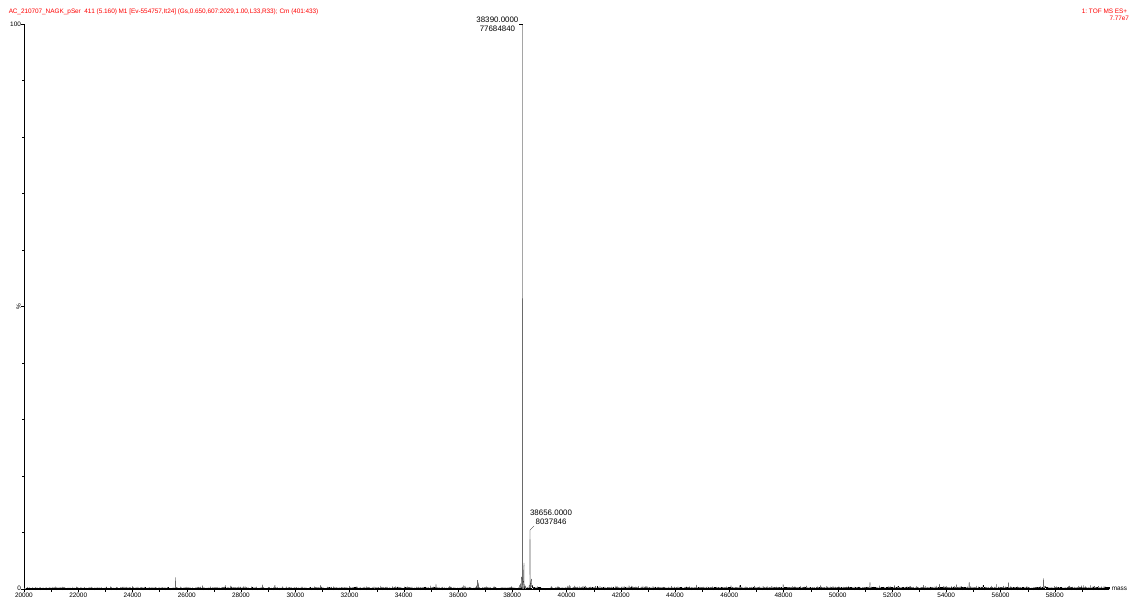

ppS76-NAGK

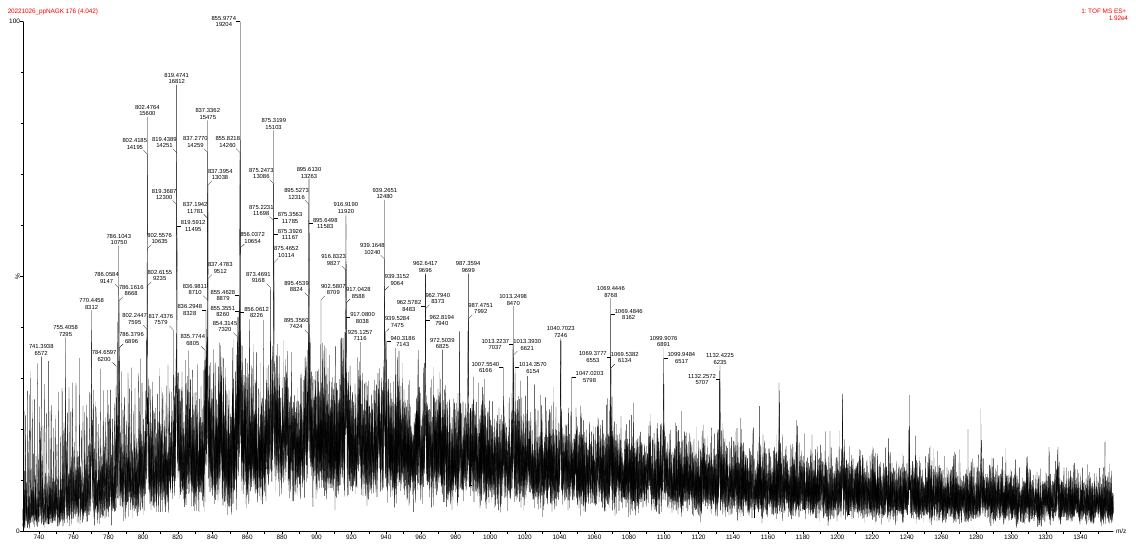

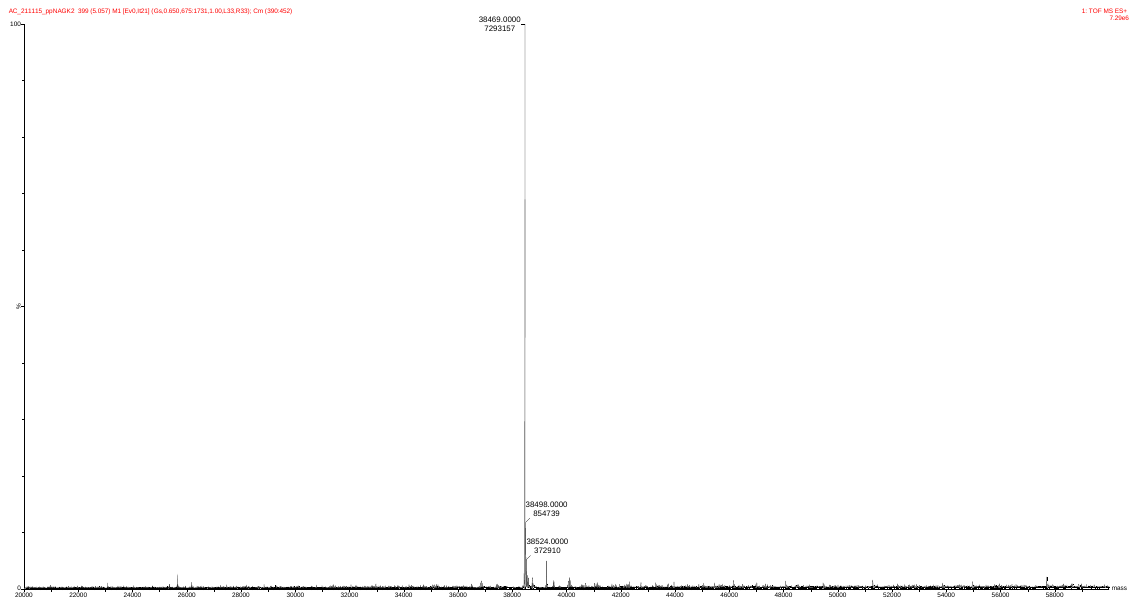

ppS76-NAGK treated with HEK293T lysate

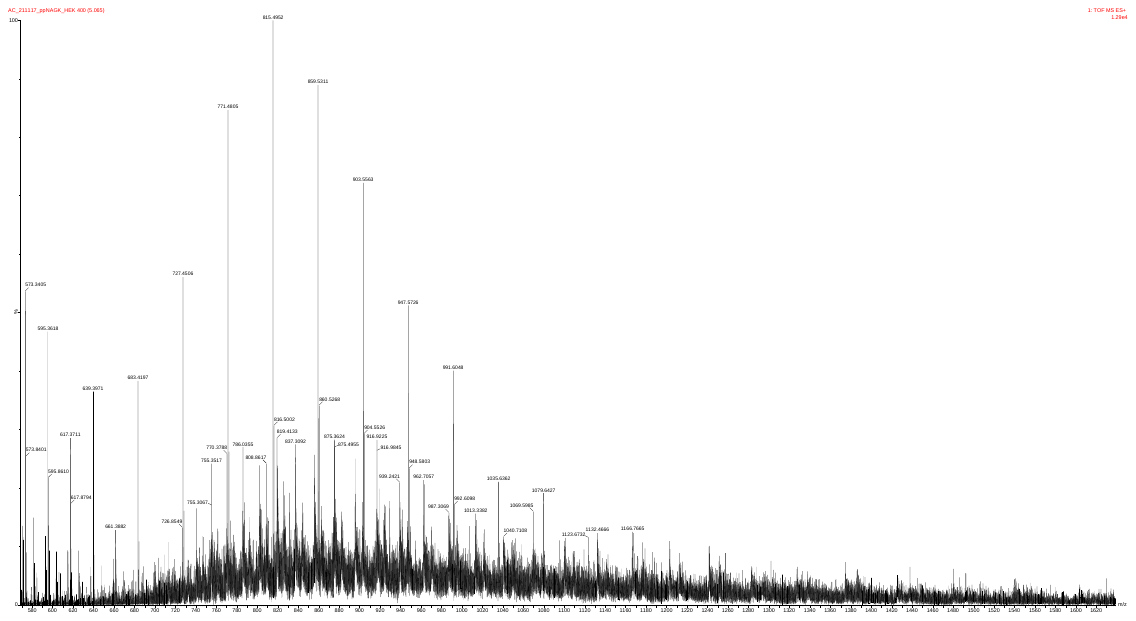

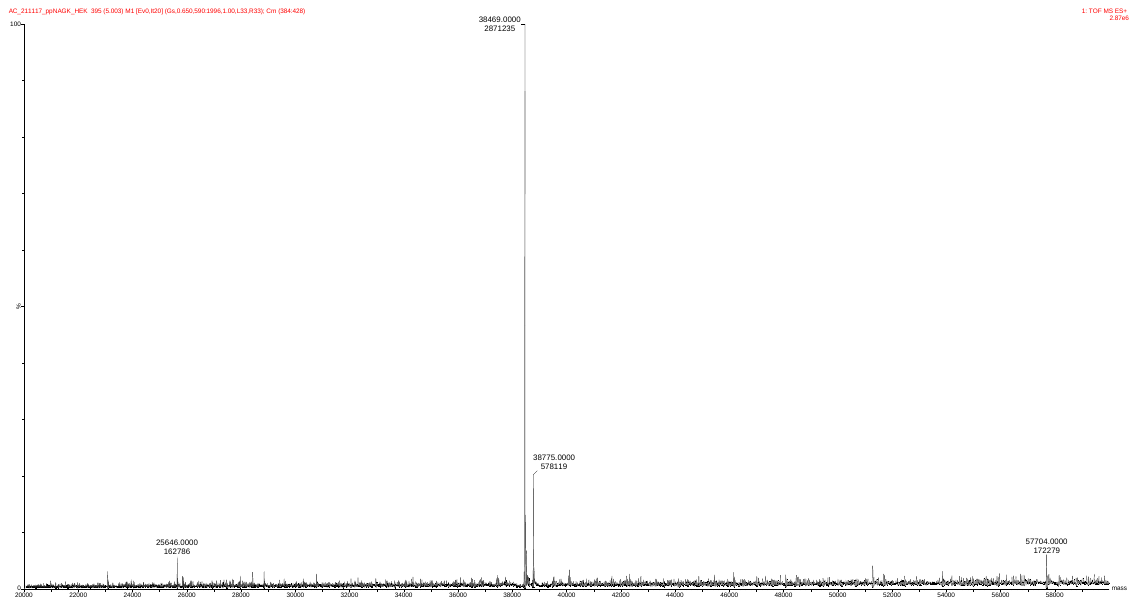

pS76-NAGK treated with HEK293T lysate

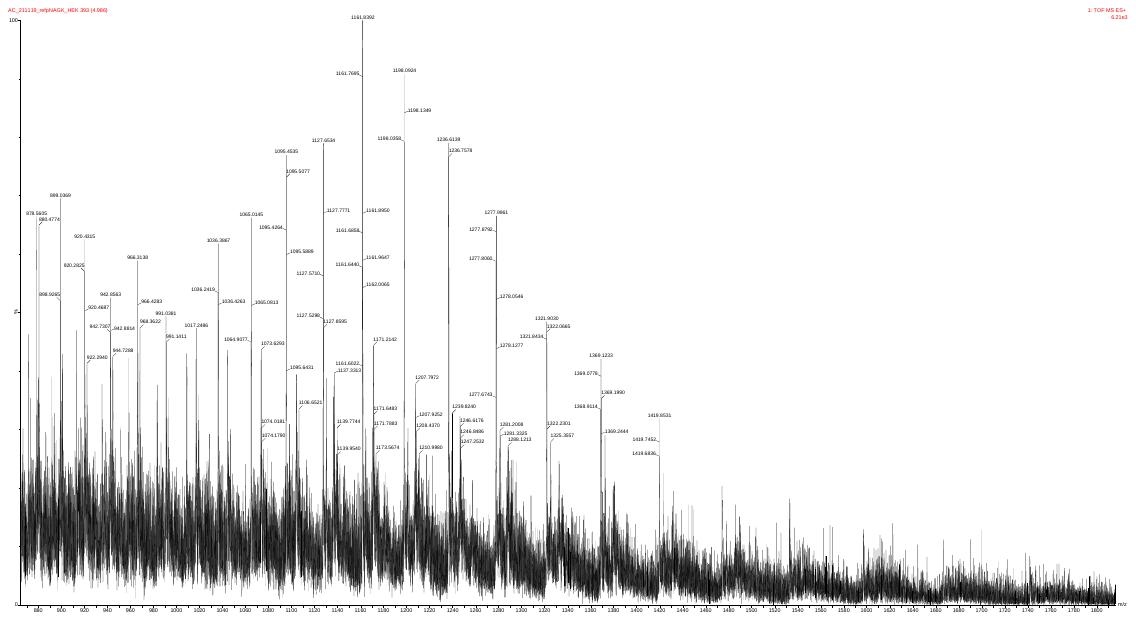

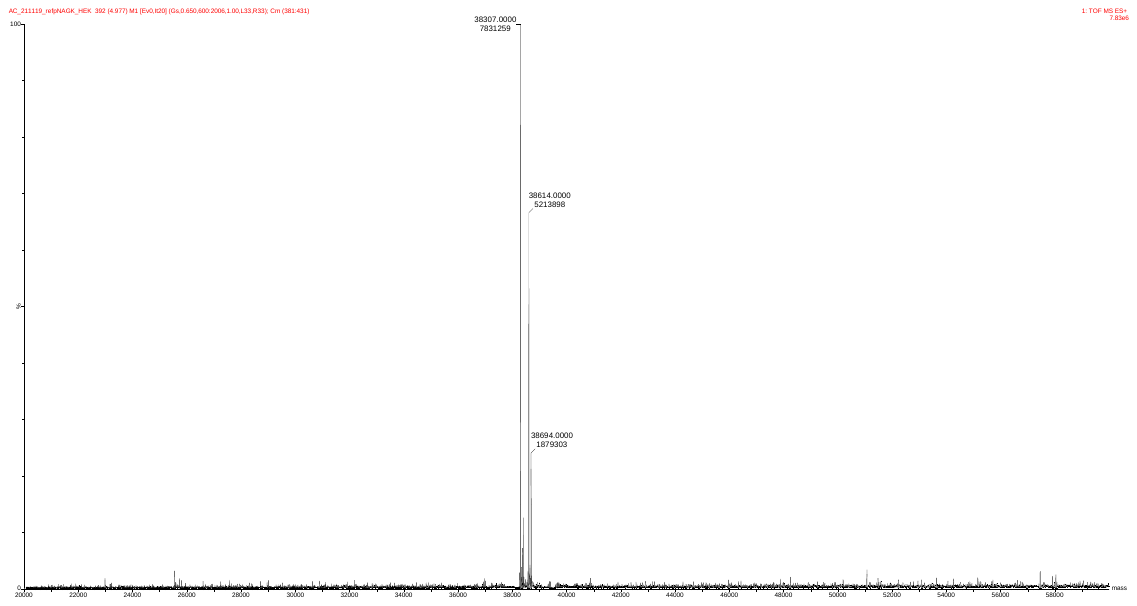

wt-NAGK treated with AurB

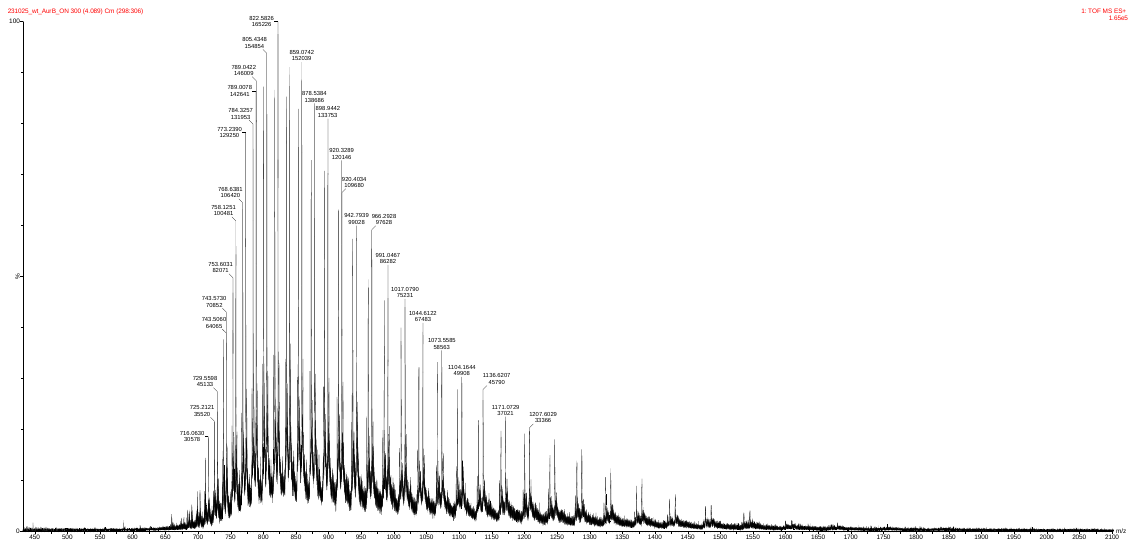

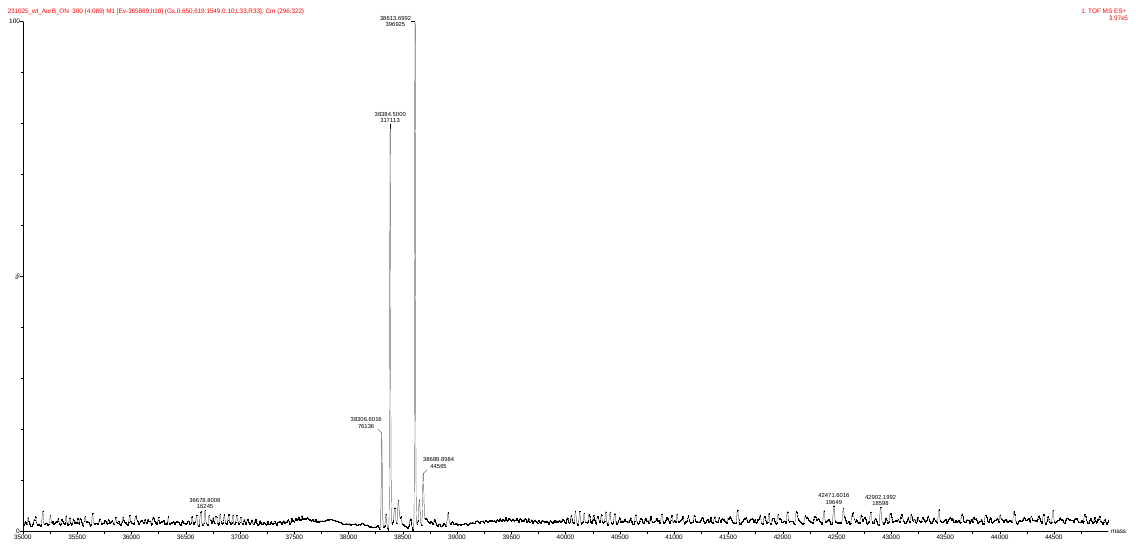

**9. Reference**

[1] [Weihofen, W.A.](https://www.rcsb.org/search?request=%7B%22query%22%3A%7B%22type%22%3A%22group%22%2C%22nodes%22%3A%5B%7B%22type%22%3A%22group%22%2C%22nodes%22%3A%5B%7B%22type%22%3A%22group%22%2C%22nodes%22%3A%5B%7B%22type%22%3A%22terminal%22%2C%22service%22%3A%22text%22%2C%22parameters%22%3A%7B%22attribute%22%3A%22audit_author.name%22%2C%22operator%22%3A%22exact_match%22%2C%22value%22%3A%22Weihofen%2C%20W.A.%22%7D%7D%5D%2C%22logical_operator%22%3A%22and%22%7D%5D%2C%22logical_operator%22%3A%22and%22%2C%22label%22%3A%22text%22%7D%5D%2C%22logical_operator%22%3A%22and%22%7D%2C%22return_type%22%3A%22entry%22%2C%22request_options%22%3A%7B%7D%7D), [Berger, M.](https://www.rcsb.org/search?request=%7B%22query%22%3A%7B%22type%22%3A%22group%22%2C%22nodes%22%3A%5B%7B%22type%22%3A%22group%22%2C%22nodes%22%3A%5B%7B%22type%22%3A%22group%22%2C%22nodes%22%3A%5B%7B%22type%22%3A%22terminal%22%2C%22service%22%3A%22text%22%2C%22parameters%22%3A%7B%22attribute%22%3A%22audit_author.name%22%2C%22operator%22%3A%22exact_match%22%2C%22value%22%3A%22Berger%2C%20M.%22%7D%7D%5D%2C%22logical_operator%22%3A%22and%22%7D%5D%2C%22logical_operator%22%3A%22and%22%2C%22label%22%3A%22text%22%7D%5D%2C%22logical_operator%22%3A%22and%22%7D%2C%22return_type%22%3A%22entry%22%2C%22request_options%22%3A%7B%7D%7D), [Chen, H.](https://www.rcsb.org/search?request=%7B%22query%22%3A%7B%22type%22%3A%22group%22%2C%22nodes%22%3A%5B%7B%22type%22%3A%22group%22%2C%22nodes%22%3A%5B%7B%22type%22%3A%22group%22%2C%22nodes%22%3A%5B%7B%22type%22%3A%22terminal%22%2C%22service%22%3A%22text%22%2C%22parameters%22%3A%7B%22attribute%22%3A%22audit_author.name%22%2C%22operator%22%3A%22exact_match%22%2C%22value%22%3A%22Chen%2C%20H.%22%7D%7D%5D%2C%22logical_operator%22%3A%22and%22%7D%5D%2C%22logical_operator%22%3A%22and%22%2C%22label%22%3A%22text%22%7D%5D%2C%22logical_operator%22%3A%22and%22%7D%2C%22return_type%22%3A%22entry%22%2C%22request_options%22%3A%7B%7D%7D), [Saenger, W.](https://www.rcsb.org/search?request=%7B%22query%22%3A%7B%22type%22%3A%22group%22%2C%22nodes%22%3A%5B%7B%22type%22%3A%22group%22%2C%22nodes%22%3A%5B%7B%22type%22%3A%22group%22%2C%22nodes%22%3A%5B%7B%22type%22%3A%22terminal%22%2C%22service%22%3A%22text%22%2C%22parameters%22%3A%7B%22attribute%22%3A%22audit_author.name%22%2C%22operator%22%3A%22exact_match%22%2C%22value%22%3A%22Saenger%2C%20W.%22%7D%7D%5D%2C%22logical_operator%22%3A%22and%22%7D%5D%2C%22logical_operator%22%3A%22and%22%2C%22label%22%3A%22text%22%7D%5D%2C%22logical_operator%22%3A%22and%22%7D%2C%22return_type%22%3A%22entry%22%2C%22request_options%22%3A%7B%7D%7D), [Hinderlich, S.](https://www.rcsb.org/search?request=%7B%22query%22%3A%7B%22type%22%3A%22group%22%2C%22nodes%22%3A%5B%7B%22type%22%3A%22group%22%2C%22nodes%22%3A%5B%7B%22type%22%3A%22group%22%2C%22nodes%22%3A%5B%7B%22type%22%3A%22terminal%22%2C%22service%22%3A%22text%22%2C%22parameters%22%3A%7B%22attribute%22%3A%22audit_author.name%22%2C%22operator%22%3A%22exact_match%22%2C%22value%22%3A%22Hinderlich%2C%20S.%22%7D%7D%5D%2C%22logical_operator%22%3A%22and%22%7D%5D%2C%22logical_operator%22%3A%22and%22%2C%22label%22%3A%22text%22%7D%5D%2C%22logical_operator%22%3A%22and%22%7D%2C%22return_type%22%3A%22entry%22%2C%22request_options%22%3A%7B%7D%7D), J Mol Biol, **2006**, 364**:** 388.
